# Phylogeny not mimicry drives chemical variation in tropical aposematic butterflies

**DOI:** 10.1101/2023.12.22.572968

**Authors:** Rachel Blow, Stephanie Ehlers, Ian Warren, Daiane Szczerbowski, Stefan Schulz, Chris Jiggins

## Abstract

Chemical signals serve a vast array of functions for both intra- and interspecific communication and defence. Chemically defended, aposematic insects will advertise their noxiousness to predators with warning signals and it has been suggested that volatile compounds shared between species could send a common signal to deter avian predators, similar to colour pattern mimicry. This is exemplified in the bright, mimetic warning colours of toxic heliconiine and ithomiine butterflies. Besides defence, males in these groups also use chemicals to communicate with potential female mates and male competitors. Here, we characterise chemical profiles of male Heliconiini and Ithomiini species from a range of mimicry rings. We find that chemical distances between species are not associated with mimicry and instead show a strong phylogenetic signal. In addition, we show that the distribution of specific shared compounds does not correlate with mimicry. We propose chemical signals in these groups are primarily used for intraspecific communication and highlight the potential selective mechanisms driving chemical profile evolution, such as selection for species-specific pheromone concentrations and collective male deterrent compounds. Our findings imply that chemical profile variation among butterfly species is the product of shared ancestry. However, more research is required on the behavioural significance of compounds among butterflies and predator perception of olfactory warning signals to identify which selective pressures are exerted on specific chemical traits.

## Introduction

Chemical signals are used to transmit a diverse array of information both within and between species. Insect pheromones are used for communication between members of the same species and are known to mediate many interactions including mating, aggregation, kin recognition and alarm call behaviours (Birch & Haynes, 1982; Wyatt, 2014; Yew & Chung, 2015). In addition, many insects use chemicals in defence, and protect themselves against natural enemies using allomones in the form of repellents, irritants, and toxins (Eisner, 1970; Pasteels et al., 1983; Ruxton et al., 2018; Sugiura, 2020). Consequently, the chemical profile of an insect can be complex in both composition and function and is exposed to a variety of selection pressures.

Aposematism is the combination of an unprofitable secondary defence with a warning signal that indicates its presence (Mappes et al., 2005; Ruxton et al., 2018). This arises through selection from predation pressure and is most often the pairing of conspicuous colouration with a noxious chemical substance, such as seen in bees and wasps. Aposematism can also be multimodal, so that additional behavioural, auditory or chemical signals and defences are involved (Rowe & Guilford, 1999; Rowe & Halpin, 2013). This is demonstrated in the bright colouration, chirping sounds, and malodorous secretions of great peacock moth (*Saturnia pyri*) larval defences (Bura et al., 2009) or the photic displays and strong odours of fireflies (Vencl et al., 2016). Complex multimodal defence and signalling strategies can be favoured if they target multiple predators or improve the efficacy of visual warning signals (Hebets & Papaj, 2005; Rowe & Guilford, 1999; Rowe & Halpin, 2013).

Müllerian mimicry improves the efficacy of aposematic warning signals via positive frequency dependent selection. The evolution of phenotype matching among defended prey species can enhance predator signal learning and reduce the individual costs of predator learning (Ruxton et al., 2018; Sherratt, 2008). As with aposematism, most examples of Müllerian mimicry describe convergence in brightly coloured visual warning signals between toxic species, famous examples including poison dart frogs, coral snakes and tropical butterflies (Bosque et al., 2022; Jiggins, 2017; Symula et al., 2001). There is also evidence of other forms of Müllerian mimicry, including acoustic mimicry in toxic moths (O’Reilly et al., 2019), as well as multimodal mimicry, through which multiple aspects of the phenotype converge between aposematic species. This is suggested to be the case among various aposematic insects that exhibit both warning colouration and warning odours (Bonacci et al., 2008; Guilford et al., 1987; Moore et al., 1990; Rothschild, 1961). As well as targeting multiple enemies or heightening the effect of visual signals, convergence of warning signals in Müllerian mimics may make aposematic individuals more distinct from their palatable Batesian mimics (Rowe & Halpin, 2013).

Unlike convergent warning signals that are shared between multiple species, pheromones tend to be species-specific. This is particularly true for sex pheromones, as they are tightly linked to mate choice (Gomez-Diaz & Benton, 2013; Symonds & Elgar, 2008; Wyatt, 2014). Due to selection for optimal mutual recognition between signaller and receiver, sex pheromones are generally thought to be under stabilizing selection (Groot et al., 2016). However, there is also growing evidence for the role of divergent selection in pheromone evolution leading to species differences (Smadja & Butlin, 2009; Symonds & Elgar, 2008). In particular, patterns consistent with reproductive isolation via reproductive character displacement and reinforcement have frequently been identified (Bacquet et al., 2015; Cama et al., 2022; Groot et al., 2006; Löfstedt et al., 1991; Shahandeh et al., 2018; Wicker-Thomas, 2011). Such divergence is likely to be particularly rapid and pronounced between closely-related, sympatric species with visually ambiguous mating signals, such as Müllerian mimics (C. Estrada & Jiggins, 2008; Gomez-Diaz & Benton, 2013; Mérot et al., 2013).

Butterflies in the tribes Heliconiini and Ithomiini are toxic and advertise unprofitability to predators with brightly coloured wing patterns. Furthermore, the high prevalence of Ithomiini butterflies in Neotropical forest communities has led to convergence in wing patterns via Müllerian mimicry (Beccaloni, 1997; Brower et al., 1963; Jiggins, 2017), resulting in the generation of distinct mimicry rings (Figures 1 and 2). Heliconiini and Ithomiini mimicry is multimodal through the combination of wing colour patterns, wing morphology and flight behaviour (Beccaloni, 1997; Hill, 2021; Srygley, 1999). Recently, the volatile chemical phenotype has also been suggested to play a role in mimicry and defence against predators, due to growing evidence for chemical convergence among butterflies and the role of olfaction in avian ecology (Avilés & Amo, 2018; Balthazart & Taziaux, 2009; Grieves et al., 2022; S. S. Steiger et al., 2008).

**Figure 1.**
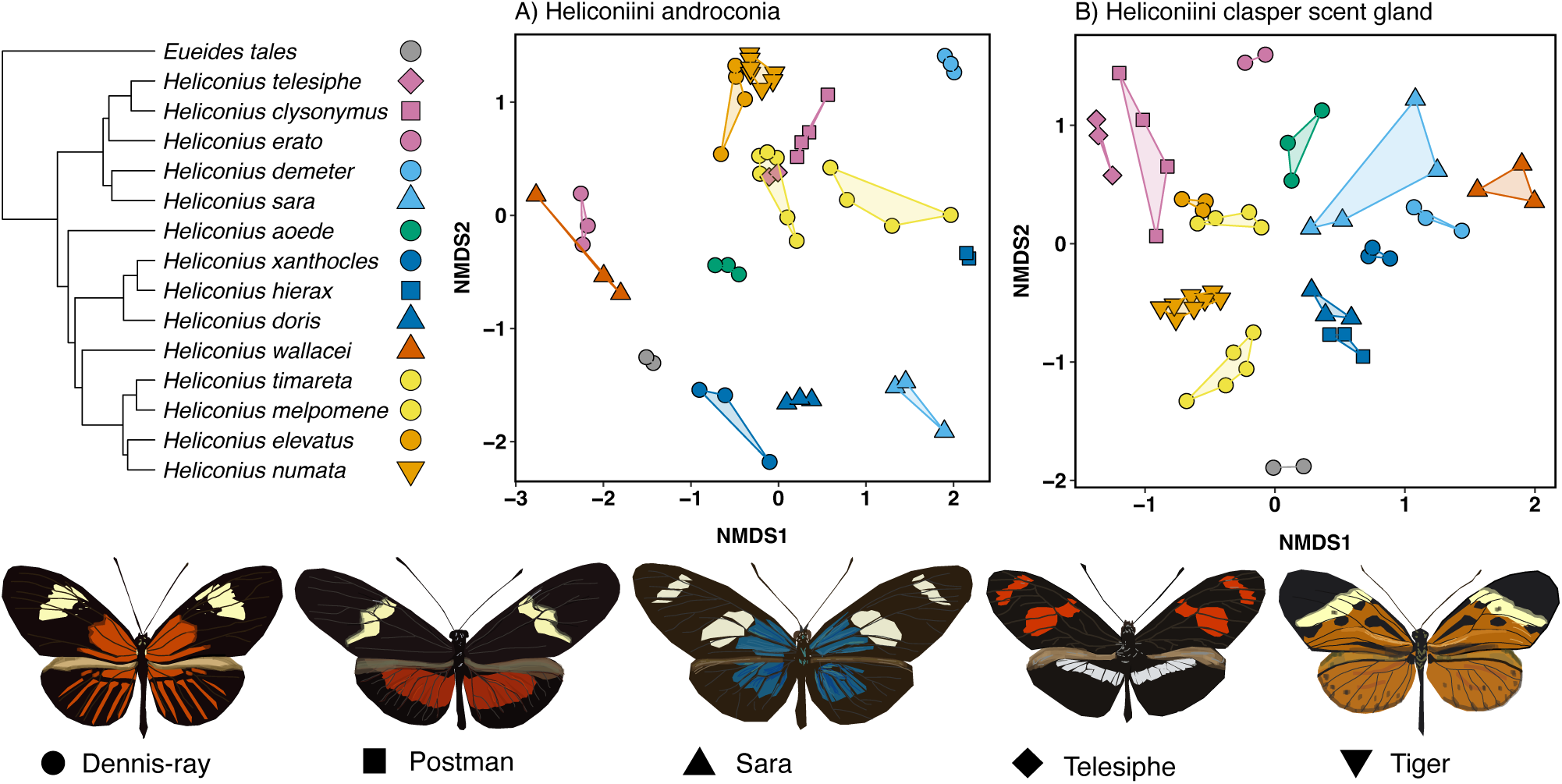
Variation in phylogeny, mimicry and chemistry among Heliconiini species. NMDS plots in the top right illustrate in two dimensions the variation in chemical distances among male androconial (A: stress = 0.16) and CSG (B: stress = 0.24) profiles. Each point represents an individual, joined to other members of the same species with lines and shaded polygons. Colours represent monophyletic clades, as outlined by colours at tips of the phylogenetic tree in the top left. Shapes represent mimicry rings, and mimicry ring classifications for each species are denoted by shapes at tips of the phylogenetic tree. Representative appearances of each mimicry ring are displayed in the legend at the bottom of the figure.

**Figure 2.**
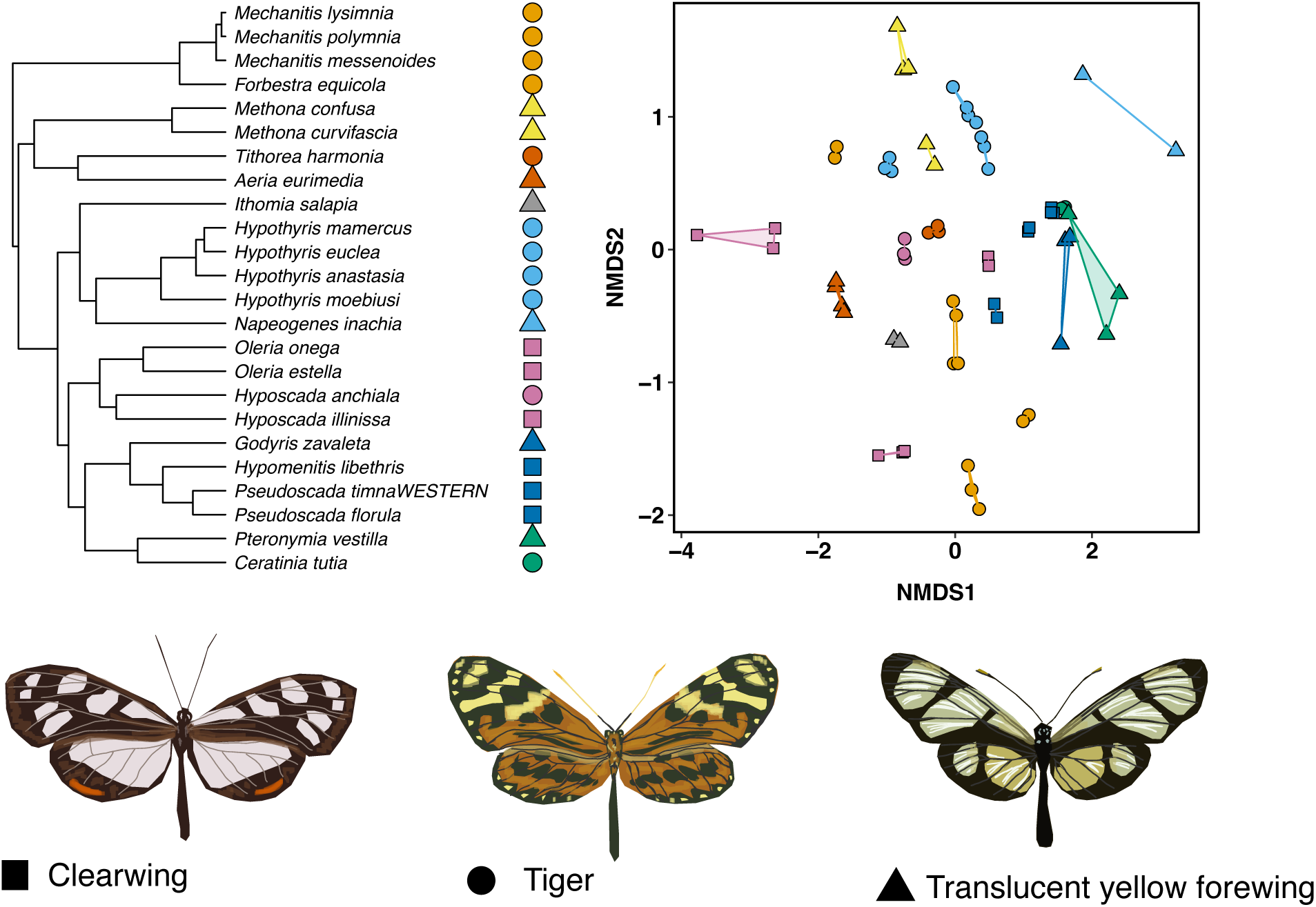
Variation in phylogeny, mimicry and chemistry among Ithomiini species. NMDS plot in the top right illustrates in two dimension variation in chemical distances among male androconial profiles (stress = 0.08). Each point represents an individual, joined to other members of the same species with lines and shaded polygons. Colours represent monophyletic clades, as outlined by colours at tips of the phylogenetic tree in the top left. Shapes represent mimicry groups, and mimicry group classifications for each species are denoted by shapes at tips of the phylogenetic tree. Representative appearances of each mimicry group are displayed in the legend at the bottom of the figure.

Chemical extracts of Ithomiini and Heliconiini butterflies are complex mixtures of compounds that contain volatiles with diverse functions (Darragh et al., 2020; Ehlers & Schulz, 2022; Stefan Schulz et al., 2008). It is possible that odours in Heliconiini and Ithomiini act as warning signals and covary with mimetic visual warning signals. In support of this, a phylogenetic comparison of Ithomiini pheromone components found no relationship between phylogeny and pyrrolizidine alkaloid derivative occurrence (Stefan Schulz et al., 2004), suggesting ecological factors, rather than phylogenetic relatedness, are driving variation in chemistry. In addition, multiple studies have detected high levels of chemical profile similarity between distantly related, mimetic species of *Heliconius* (Darragh et al., 2020; Catalina Estrada et al., 2011; Mann et al., 2017), as well as high differentiation between mimetic forms of *H. erato* (Ehlers et al., 2021). Recent research has revealed the importance for the use of smell in bird foraging, reproduction and risk assessment (Avilés & Amo, 2018). Because avian predators drive colour pattern mimicry in these butterflies (Langham, 2004, 2006; Pinheiro, 1996, 2003), predation pressure from birds may also play a role in shaping chemical variation among populations.

Ithomiini and Heliconiini volatiles also function as pheromones. Male Ithomiini use chemical signals in female mate attraction, competitor repellence and aggregation. Males gather pyrrolizidine alkaloids from wilting Solanaceae plants for use as pheromone precursors (Boppré, 1986; Edgar et al., 1976; Eisner & Meinwald, 1987; Pinheiro et al., 2008; Pliske, 1975; Stefan Schulz et al., 2004). Chemical signalling also plays an important communicative role in *Heliconius* and pheromones are a mix of biosynthesized compounds and volatiles that are transformed or obtained directly from larval host plant compounds. Males have specialized hindwing androconia, from which pheromones involved in female mate choice are released (Byers et al., 2020; Darragh et al., 2017; González-Rojas et al., 2020). In addition, males transfer antiaphrodisiac pheromones from clasper scent glands (CSG) to females during mating, where they deter subsequent male mating attempts (Gilbert, 1976; Melo et al., 2022; Stefan Schulz et al., 2008). Female-released pheromones are also thought to be involved in species recognition by males, but less is known about this in terms of chemistry and behaviour (C. Estrada & Jiggins, 2008; Jiggins et al., 2004; Mérot et al., 2015).

Here, we sampled sympatric species from a single geographic region, to compare the chemical extracts of males from 15 and 25 species of Heliconiini and Ithomiini, respectively. The species studied vary in phylogenetic relatedness and represent several distinct mimicry rings (Figures 1 and 2). Previous research suggests chemical signals may converge between aposematic and visually mimetic butterfly species, due to selection from avian predation pressure. However, there is also a clear role for pheromones and mate choice in shaping Heliconiini and Ithomiini chemistry. This raises the question: which selective pressures have been most influential in shaping semiochemical evolution in aposematic and mimetic insects? And does mimicry or phylogeny have a greater impact on chemical profile? To answer these questions, we identified macroevolutionary patterns in butterfly volatile chemistry and measured the strength of associations between chemical, mimetic and phylogenetic distances.

## Methods

### Fieldwork

We caught wild Heliconiini and Ithomiini male butterflies with hand nets during March 2019 and made provisional species identifications in the field, supplemented with subsequent DNA barcoding. We dissected Heliconiini androconial tissue from the hindwing, Heliconiini CSG tissue from the tip of the abdomen, and Ithomiini androconial hairpencils from the hindwing. For volatile extraction and internal standard calibration, we saturated dissected tissues in 200 μl dichloromethane containing 200 ng tetradecane-2-yl-acetate for one hour. For subsequent species identification and mimicry ring classification, we preserved butterfly bodies in ethanol (>99%) and dissected and photographed forewings.

### Chemical analysis

We analysed extracts by gas chromatography and mass spectrometry (GCMS) using an Agilent model 5977 mass-selective detector connected to an Agilent GC model 7890B and equipped with an Agilent ALS 7693 autosampler using a HP-5MS fused silica capillary columns (Agilent, 30 m × 0.25 mm, 0.25 µm). Injection was performed in splitless mode (250°C injector temperature) with helium as the carrier gas (constant flow of 1.2 ml/min). The temperature program started at 50°C, was held for 5 min, and then rose at a rate of 5°C/min to 320°C, before being held at 320°C for 5 min. We identified components by comparison of mass spectra and retention indices against a custom reference library in AMDIS (Mallard & Reed, 1997) and quantified using the internal standard area. To increase the signal to noise ratio, we removed any peaks with a ratio score < 50 for Heliconiini or <20 for Ithomiini samples. We similarly removed heavy compounds eluting after hexacosane for androconial extracts and nonacosane for CSG extracts (Darragh et al., 2017), as well as all compounds that were not found in at least two individuals of a given species. To reduce the amount of variation caused by life history and physiological state differences between individuals, we normalised data by dividing the concentration of each compound by the total concentration of compounds detected per individual. Finally, we removed all known contaminants.

### COI barcoding for species identification

We extracted and purified high molecular weight DNA with magnetic beads as described in: dx.doi.org/10.17504/protocols.io.b46bqzan. We amplified a 376 bp region of mitochondrial COI using the primers listed in Table S4, which contain target sites for i5- and i7-index primers for Illumina sequencing. A second round of PCR added unique combinations of i7- and i5-index barcodes to each PCR product. We pooled and cleaned barcoded PCR products using homemade beads (recipe can be found in: dx.doi.org/10.17504/protocols.io.b46bqzan). 250bp paired-end sequencing on a NovaSeq PE250 was carried out by Novogene (UK). We processed reads using the R package “ShortRead”, version 1.54.0 (Morgan et al., 2009). We then compared reference sequences against a library of 39, 354 nymphalid butterfly COI sequences, representing 3679 species, downloaded from GenBank (Benson et al., 2013), using the Basic Local Alignment Search Tool, version 2.4.0 (Altschul et al., 1990). Sequences were deposited in the European Nucleotide Archive (ENA) with project accession number PRJEB69300.

### Species identification and mimicry ring classification

We identified species by morphology using field identifications and forewing photographs. Where possible, we complemented this with COI barcodes and pre-existing genomic data. Following identification, we categorised Heliconiini species into mimicry rings and Ithomiini species into mimicry rings and broader mimicry groups. Mimicry ring designations were based on overall phenotypic similarity, the presence or absence of wing colour pattern elements, and species ecology. Ithomiini mimicry groups incorporate mimicry rings with broadly similar patterns.

### Statistical analysis

Due to the multivariate and zero-biased nature of chemical data, we used non-metric multidimensional scaling (NMDS) to visualise chemical profile distances among sampled individuals. We performed ordinations in two dimensions using a Bray-Curtis dissimilarity index with the “metaMDS” function in the package *vegan*, version 2.6.4 (Oksanen et al., 2022). To determine which factors influence profile variation, we first assessed the marginal effects of species, clade and mimicry on individual chemical profile using permutational multivariate analysis of variance (PERMANOVA) with the “adonis2” function in *vegan*. Our first hypothesis predicts that chemical profiles should vary with visual mimicry due to selection for multimodal Müllerian mimicry. To test whether chemical distances between species were significantly associated with mimicry, we calculated typical species chemical profiles by averaging all non-zero compound concentrations across sampled individuals per species. Next, we calculated species chemical distance scores using the Bray-Curtis dissimilarity index in the “vegdist” function in *vegan*. We calculated mimetic distance matrices by assigning comparisons between mimetic species pairs a score of zero, and non-mimetic species pairs a score of one. Finally, we calculated phylogenetic distance matrices using the “cophenetic.phylo” function in the package *ape,* version 5.7.1 (Paradis & Schliep, 2019) using published Heliconiini (Kozak et al., 2015) and Ithomiini (Chazot et al., 2019) phylogenies. We then tested for correlations between species chemical and mimetic distance matrices, while controlling for phylogenetic distance, using partial Mantel tests in *vegan*. Our alternative hypothesis expects the chemical profiles of more closely related species to resemble each other due to shared ancestry. We measured the amount of phylogenetic signal among species chemical profiles using Mantel correlations between species chemical and phylogenetic distance matrices, in *vegan*.

Rather than mimicking the full chemical profile, it is possible that only subsets of key compounds are used in chemical mimicry. To determine whether any individual compounds are associated to particular mimicry rings, we used the “multipatt” function in the package *indicspecies*. Compounds with a combined *specificity* (found only in that group) and *fidelity* (found in all members of that group) statistic of 0.95 or higher were accepted. For the Heliconiini androconial and CSG compounds, associations were tested among mimicry rings. For Ithomiini androconial compounds, associations were first tested among mimicry rings, then broader mimicry groups.

Selection for reproductive isolation could, similarly, drive the evolution of a few key species-specific recognition compounds. We used the same function to identify compounds with specificity and fidelity to individual species. All statistical analyses were performed in *R* version 4.2.3 (R Core Team, 2023) and all figures were plotted using the package *ggplot* version 3.4.1 (Wickham, 2016).

## Results

Heliconiini and Ithomiini butterflies are found throughout South and Central America and exhibit striking subspecies variation. To reduce the influence of factors such as host plant availability and geographic isolation on chemical diversity, we collected sympatric subspecies from the rainforest and Andean foothills surrounding Tena, Ecuador. Extracts were collected from 15 species of Heliconiini and 23 species of Ithomiini. The Heliconiini were distributed among five mimicry rings, (Figure 1; dennis-ray, postman, sara, telesiphe, and tiger). Ithomiini species were distributed among nine mimicry rings (agnosia, aureliana, confusa, eurimedia, hermias, lerida, libethris, mamercus, and mothone), which were in turn distributed among three broader mimicry groups (Figure 2; clearwing, tiger, and translucent yellow forewing). Of the Heliconiini, 14 species were in the genus *Heliconius* and one was from the sister genus *Eueides* (Figure 1). The Ithomiini species collected spanned 15 genera (Figure 2). The number of individuals per species ranged from two to eight. Two different subspecies of *H. numata* were collected, but as these were in the same mimicry ring and chemical variation did not segregate by subspecies, all individuals were treated as one.

In total, 109 androconial and 443 CSG compounds were found in extracts from 54 Heliconiini males. There was considerable variation between individuals (Figure 1), with species significantly predicting marginal variation in both androconial (PERMANOVA, species, F_3,53_ = 16.74, *p < 0.001*) and CSG (PERMANOVA, species, F_3,53_ = 9.54, *p < 0.001*) chemical profiles. The average number of compounds per species in androconial extracts ranged from two in *H. sara* to 33 in *H. clysonymus* (Figure S1A), and in CSG extracts from 15 in *H. sara* to 79 in *H. timareta* (Figure S1B).

The average total amount of compound per species ranged from 69 ng in *H. sara* to 28,032 ng in *H. demeter* for androconial compounds (Figure S1A) and from 2027 ng in *H. sara* to 162,816 ng in *H. elevatus* for CSG compounds (Figure S1B). In species with high total amounts of androconial and CSG compounds, this was driven by the large abundance of a few major compounds (Figures S3 and S4), as is characteristic of *Heliconius* (Darragh et al., 2020; Catalina Estrada et al., 2011; Stefan Schulz et al., 2008).

We found 143 androconial compounds in extracts from 63 Ithomiini males. Chemical profiles varied considerably between individuals (Figure 2) with species significantly predicting variation (PERMANOVA, species, F_8,62_ = 10.82, *p < 0.001*). The average number of compounds per species ranged from two in *Oleria onega* to 41 in *Tithorea harmonia* (Figure S2A). The average total amount of compound per species ranged from 72 ng in *Oleria estella* to 24,239 ng in *T. harmonia* (Figure S2B). Like Heliconiini chemical profiles, Ithomiini androconial extracts are dominated by a small number of very abundant compounds and additional minor compounds.

We detected strong phylogenetic signal in Heliconiini chemical extracts, as phylogenetic distance correlated with both androconial chemical distance (Mantel test, r = 0.25, *p < 0.05*) and CSG chemical distance (Mantel test, r = 0.4, *p* < 0.001) (Figures 3A and B). However, when controlling for phylogenetic relatedness, mimetic distance did not correlate with Heliconiini androconial chemical distance (partial Mantel test, r = -0.07, *p* = 0.73) or CSG chemical distance (partial Mantel test, r = 0.02, *P* = 0.4).

**Figure 3.**
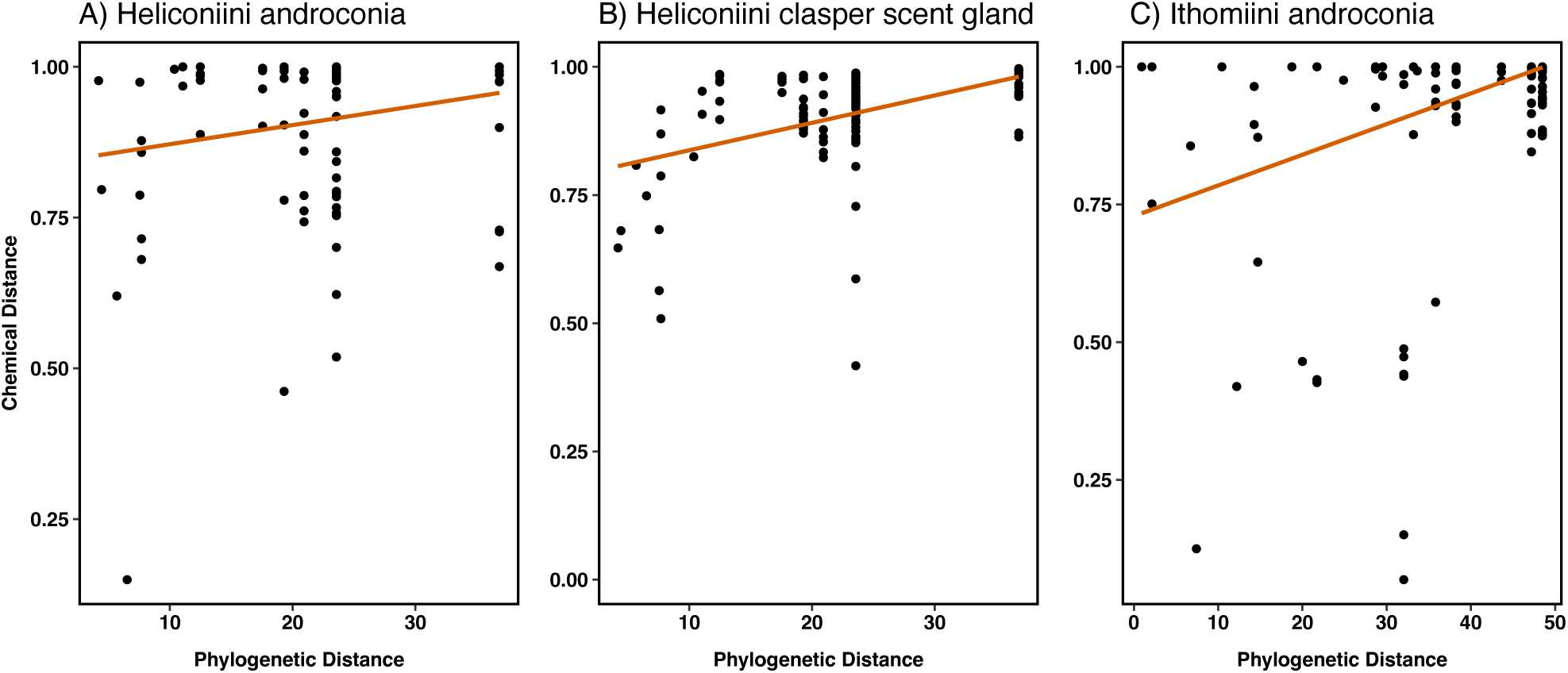
Phylogenetic distance versus Heliconiini androconial (A), Heliconiini CSG (B), and Ithomiini androconial (C) chemical distances. Points indicate pairwise distance scores between species. Red regression lines are used here to represent significant Mantel test correlations.

We identified several Heliconiini androconial and CSG compounds shared among multiple species, though no individual compounds were significantly associated with a mimicry ring. Of the androconial compounds: pentacosane, heneicosane, tricosane and syringaldehyde are shared by 50% of the sampled species (Figure S3). The former compound is found in species from all mimicry rings. The latter three compounds are found in species from all mimicry rings excluding sara. Of the CSG compounds: 18 compounds are shared by 50% of the sampled species (Figure S4). Heptacosane is present in all species and 1,3-docosanediol is present in 80% of species from all mimicry rings. Other notable shared compounds include (*E*)-β-ocimene, which is a known *Heliconius* antiaphrodisiac pheromone (Stefan Schulz et al., 2008) and an unidentified compound (here named unknown B135) that has previously been shown to be highly electrophysiologically active (unpublished data) in *Heliconius*. Both are shared by 50% of the species sampled here, with (*E*)-β-ocimene present in species from all mimicry rings except tiger and the unidentified GC-EAD active compound present in species from the dennis-ray and sara mimicry rings only.

We identified candidate Heliconiini androconial recognition compounds that had high specificity and fidelity to a given species (Table S1). Some of these compounds are known to play a pheromonal role in *Heliconius* and other insects. For example, linalool oxide (furanoid), a suggested indicator compound of *H. xanthocles*, is used by *H. charithonia* males to find and identify immature, conspecific females (Catalina Estrada et al., 2010), and nonanoic acid, the suggested indicator compound of *H. clysonymus*, is used as a pheromone by multiple insect species (El-Sayed, 2023). For five species, multiple compounds were significant indicators of species identity, such as *H. demeter*, which has 11 significantly species-specific compounds (Table S1). Five candidate recognition compounds were proposed for the species *H. heirax*, one of which is the aliphatic nitrile octadecanenitrile. To the best of our knowledge, this is the first aliphatic nitrile reported from insects. For four species, the suggested indicator compounds did not pass the combined statistic threshold of 0.95.

We also identified several species-specific CSG compounds (Table S2). Again, some of these compounds have a known pheromonal role in insects, such as ethyl oleate, which is one of the suggested indicator compounds for *H. timareta*, and functions as a primer pheromone in honey bees (Leoncini et al., 2004). Others have a known role in defence, such as 2-sec-Butyl-3-methoxypyrazine, also found in *H. timareta* (Burdfield-Steel et al., 2018). However, many of the species-specific CSG compounds have not yet been functionally characterized, and for some the structure is also yet to be identified.

Ithomiini androconial extracts exhibited phylogenetic signal, and phylogenetic distance correlated with androconial chemical distance (Figure 3C; Mantel test, r = 0.4, *p* < 0.001). When controlling for phylogenetic relatedness, Ithomiini mimetic distance did not correlate with androconial chemical distance by mimicry ring (partial Mantel test, r = 0.02, *p* = 0.28) or mimicry group (partial Mantel test, r = -0.04, *p* = 0.71).

The androconial compound 4-hydroxyphenylethanol was significantly associated with the confusa mimicry ring (multilevel pattern analysis, combined statistic = 1, *p* < 0.05), part of the translucent yellow forewing mimicry group. However, this mimicry ring contains only two closely related and almost identical species in the genus *Methona*. It is therefore possible that this effect is driven by either phylogenetic relatedness or mimicry, or both. No Ithomiini androconial compounds were significantly associated with any mimicry group.

Certain Ithomiini androconial compounds are shared by multiple species, unrelated to mimicry (Figure S5). The most common compounds are pentacosane, shared by 30% of species, along with ithomiolide C, oxodanaidal, and an unidentified compound (here named unknown B104), which are shared by 20% of species (Figure S5). Pentacosane and ithomiolideC are present in species in all mimicry groups, but not all mimicry rings. Oxodanaidal and the unidentified compound B104 are unique to the tiger mimicry group but are found in less than 50% of species in this group.

We identified candidate Ithomiini species recognition compounds that had significant specificity and fidelity to individual species (Table S3). Like the Heliconiini candidate indicator compounds, this included compounds with known pheromonal functions in other insects. For example, 1-hexadecanol, a suggested indicator of *Hypothyris moebuisi* and female sex pheromone in *Helicoverpa armigera* (Zhang et al., 2012), and octadecanal, significantly specific to *Hypothyris euclea*, and used as a courtship signal in *Heliconius melpomene* (Byers et al., 2020). For many species, multiple indicator compounds are suggested, such as *Tithorea harmonia,* which has 18 compounds with significant species specificity and fidelity (Table S3). The suggested indicator compound of one species, *Pseudoscada florula*, did not meet the 0.95 combined statistic requirement, and no indicator compounds were detected for *Hypothyris mamercus*.

## Discussion

Our results show that male androconial and clasper scent gland (CSG) chemical profiles of tropical butterflies contain complex blends of compounds that vary both qualitatively and quantitatively among species. This was expected from previous descriptions of complex Heliconiini and Ithomiini butterfly scent chemistry (Cama et al., 2022; Darragh et al., 2017, 2020; Ehlers et al., 2021; Mann et al., 2017; S Schulz et al., 1988; Stefan Schulz et al., 2004). Chemical blends are species-specific and exhibit varying levels of dissimilarity between species, dependent on phylogenetic relatedness. Variation in blend is not associated with mimicry in Heliconiini androconial or CSG chemical profiles, nor Ithomiini androconial chemical profile. Similarly, though several compounds are shared between species within both butterfly groups, this cannot be explained by mimicry.

### Chemistry is not associated with mimicry

The occurrence of multiple sympatric mimicry rings among Heliconiini and Ithomiini is thought to be maintained in part by differentiation in visual signal efficacy across microhabitats (Beccaloni, 1997; Elias et al., 2008; Jiggins, 2017; Mallet & Gilbert, 1995). It is possible that chemical signals do not converge with visual signals because they do not vary across microhabitats in the same way. For example, Neotropical forest microhabitats have different avian predator community assemblages that also likely have different colour signal detection biases (Jiggins, 2017; Rowe & Skelhorn, 2004; Willmott & Mallet, 2004). Olfactory capabilities have been shown to differ between bird species but variation is largely dependent on aquatic versus terrestrial habitats and herbivore versus carnivore diets, which are necessarily invariable among Heliconiini and Ithomiini bird predators (Avilés & Amo, 2018; Manger et al., 2015).

An alternative explanation is that predator capabilities do not vary between habitats, but that the same predators learn different mimicry rings in a context dependent manner (Jiggins, 2017), such as an aversion to the colour red associated with open, well-lit forest glades. Insects are well-known for their ability to distinguish between olfactory cues in a context-dependent manner (Alkema et al., 2019; Mitaka & Akino, 2021; Orlova & Amsalem, 2021). There is also some evidence of sex- and context-dependent use of olfaction versus vision during great tit foraging in urban and forest environments (Rubene et al., 2023).

If the perception of olfactory signals by bird predators shows little variation across microhabitats, warning signals might be shared by all prey species, regardless of visual mimicry. For instance, many aposematic species, representing a variety of brightly-coloured phenotypes, produce the same pyrazines (Moore et al., 1990), with which birds are able to form a conditioned aversion (Guilford et al., 1987). One *Heliconius* CSG compound, heptacosane, is shared by all species. This compound is known to mediate avoidance of insect predators in aphids, and is used as a trail pheromone by some ladybird species (Nakashima et al., 2006). It would be interesting to test the ability of birds to form associations between heptacosane and unprofitable prey to determine the biological role of this compound in Heliconiini butterflies.

### A defensive role for pyrazines

The CSG extracts of *H. timareta* males contain 2-sec-butyl-3-methoxypyrazine. Pyrazines have a known defensive function against predators in multiple aposematic insect species (Avery & Nelms, 1990; Burdfield-Steel et al., 2018; Guilford et al., 1987; Rowe & Guilford, 1999; José R. Trigo, 2000). These compounds have also been suggested to be involved in warning odour Müllerian mimicry (Dettner & Liepert, 1994; Rothschild, 1961). It is therefore surprising that the presence of these compounds is restricted to *H. timareta*. A recent study, describing inter- and intraspecific chemical variation in *Heliconius*, detected pyrazines in the CSG extracts of certain subspecies of *H. cydno* and *H. melpomene* (Darragh et al., 2020), however they too found that pyrazines were lacking from the chemical profiles of the equivalent Ecuadorian subspecies.

While recent evidence suggests insects are able to biosynthesize pyrazines (Burdfield-Steel et al., 2018), the lack of consistency in pyrazine presence among *Heliconius* species and subspecies suggests these compounds are obtained directly from another source. It is possible that pyrazines are sequestered from host plants specific to *H. timareta*, however *Passiflora* are not known to produce these compounds. Alternatively, pyrazine production by microbial endosymbionts is common in insects (Calcagnile et al., 2019). Because symbiotic and mutualistic interactions are contingent on the co-evolution and ecological niche specialisation of species, microbe-produced pyrazines could be restricted to certain *Heliconius* lineages and geographic regions. Male hairpencils of the danaid butterfly, *Idea leuconoe,* contain a microbe-produced compound, mellein (Nago & Matsumoto, 1994). This compound is thought to be used for defence by the butterfly (Nishida et al., 1996), which in turn serves as a vector of fungal spores. Interestingly, we detected the same compound in the CSG chemical profile of *H. erato, H. clysonymus,* and *H. doris*. These findings suggest there may be an ecological role for microbe-produced volatiles in Heliconiini communication and defence.

### Why are compounds shared between species?

The extracts of Heliconiini and Ithominiini male androconia and CSG tissue contain numerous compounds that are shared among distantly related species in what seem to be unpredictable patterns. We have shown that this cannot be readily explained by multimodal mimicry for defence against avian predators. However, it is possible that visual and chemical signalling channels are used to deter different sets of predators. For example, there is evidence that bats and spiders predate on *Heliconius* at night, when their wing colour patterns are not detectable (Salcedo, 2011). The distribution of chemical signals aimed at nocturnal predators would therefore not be predictable by visual mimicry and may explain why a number of compounds in our dataset are shared between multiple species in different mimicry rings. For example, the monoterpene (*E*)-β-ocimene is shared by seven Heliconiini species of varying degrees of relatedness found across four mimicry rings. In the defensive secretions of papilionid butterflies, monoterpenes, including (*E*)-β-ocimene, are most effective against invertebrate predators (Honda, 1981; Palma-Onetto et al., 2023), and may be used for the same purpose in *Heliconius*.

Male Ithomiini use chemical signals to communicate with both con- and heterospecific male and female butterflies. Such signals may be used for female attraction, aggregation and male-male territory defence (Boppré, 1986; Edgar et al., 1976; Eisner & Meinwald, 1987; Pinheiro et al., 2008; Pliske, 1975; Stefan Schulz et al., 2004). The shared use of PA-derived pheromones among ithomiines may be related to their function as honest signals of male quality. In the PA sequestering arctiine moth, *Utetheisa ornatrix*, females select mates based on the production of hydroxydanaidal (Eisner & Meinwald, 1987, 1995). This compound advertises both direct benefits to females, in the form of PA load for predator protection, as well as the heritable ability to sequester alkaloids in her sons (Conner & Iyengar, 2016). This is thought to have driven convergence in the use of hydroxydanaidal in many PA sequestering arctiine moths (Conner & Iyengar, 2016; Stefan Schulz, 2009), and may be the reason for shared chemistry among ithomiine species.

Shared signals between ecologically overlapping species may otherwise be selected for if they mediate territorial disputes among males of all species guarding similar territories. In addition, Ithomiini butterfly aggregations are thought to be formed as a consequence of adverse conditions during the dry season (Pinheiro et al., 2008), as heterospecific males and females aggregate at shady and humid pockets of habitat. If these pockets are infrequent enough, the benefits of responding to heterospecific aggregation signals to find suitable habitat would outweigh the costs of attracting and sharing resources with heterospecifics, so that shared aggregation signals between species are favoured. Currently, we do not know the function of specific androconial compounds in Ithomiini species. Attributing functions to volatiles will help us to understand which selective forces shape their distribution among species.

Heliconiini also use male-male chemical signalling. The monoterpene (*E*)-β-ocimene is known to be used as an antiaphrodisiac by *H. melpomene* (Stefan Schulz et al., 2008). Antiaphrodisiacs are transferred from males to females during mating and deter subsequent male mating attempts (Malouines, 2017). It is possible that (*E*)-β-ocimene has the same antiaphrodisiac function in other species. For example, this is suggested to be the case in *H. erato,* although the occurrence of this compound is not consistent among subspecies and another compound, phyllisolide, is thought to function as the main antiaphrodisiac in *H. erato phyllis* (Melo et al., 2022). Shared antiaphrodisiac signals between species could help to reduce female mating harassment by both intra- and interspecific males among partially mimetic females. If time spent chasing mates is costly, the initial attraction of males to any moving object that resembles a conspecific female (Mallet et al., 1990), may impose selection for shared olfactory deterrents. Most species in which we found (*E*)-β-ocimene have a common, prominent red feature to their aposematic wing pattern, known to initiate chasing of females by males (Crane, 1955). However, other species with the same red feature are lacking (*E*)-β-ocimene, and *H. sara*, which does contain (*E*)-β-ocimene, is blue and yellow. It is possible that the shared occurrence of this compound is therefore driven by multiple evolutionary processes, and (*E*)-β-ocimene serves different functions in different species. Insect pheromones are often complex blends of compounds, and heterospecifics may respond differently to the same compound, depending on the concentration (Kaae et al., 1973; Löfstedt et al., 1991; Symonds et al., 2011; C. Y. Yang et al., 2009) and accompanying volatiles (Alkema et al., 2019; Mitaka & Akino, 2021; Orlova & Amsalem, 2021).

### Species-specific compounds for conspecific recognition

Even though many individual compounds are shared between species, we found chemical blends to be highly species-specific, a feature that may be indicative of evolution primarily for mate recognition (Gomez-Diaz & Benton, 2013; Symonds & Elgar, 2008; Wyatt, 2014). This is further supported by the fact that many of the shared compounds detected in this study function as pheromones in other insects (Yang *et al*., 2017; Funaro *et al*., 2018; Wang *et al*., 2023). This includes octadecanal, which is used by *H. melpomene* in female mate choice (Byers et al., 2020), and is found in varying concentrations across four Heliconiini and three Ithomiini species. This compound is found in particularly high concentrations in *H. melpomene* males. However, supplementing the wings of courting males with octadecanal increases mating latency in females (Byers et al., 2020). This suggests females are sensitive to specific concentrations of octadecanal and pheromone divergence between species may be partially generated by quantitative differences in overlapping compounds (Gomez-Diaz & Benton, 2013; Smadja & Butlin, 2009). In the same study, syringaldehyde was present in similar amounts in *H. melpomene* and *H. cydno* male androconial extracts, and elicited female responses in both species (Byers et al., 2020). We found syringaldehyde in a total of seven out of 15 Heliconiini species. It would be interesting to test the behavioural significance of this compound in these species and ascertain whether it works synergistically with other compounds to elicit behavioural responses in females.

We found several compounds that are present only in one species, and in all members of that species. Such compounds may serve as species recognition pheromones (Darragh et al., 2020), though their use as pheromones needs behavioural verification. The Heliconiini species *H. timareta* and *H. erato* had twice (n=16) and four times (n=32) as many species-specific CSG compounds as the group average (n=8), respectively. The Ithomiini species *Tithorea harmonia* had four times (n=19) as many species-specific androconial compounds as the group average (n=5). Proximally, this may be the result of increased ability to biosynthesise pheromones or sequester them from larval host plants. A recent study found that the Sara/Sapho + Erato stem and Silvaniform/Melpomene clades have undergone expansions in genes linked to pheromone biosynthesis (Cicconardi et al., 2023). It is therefore possible that diversification of biosynthetic enzymes used for pheromone production has resulted in a greater diversity of products in *H. erato* and *H. timareta* (Gomez-Diaz & Benton, 2013; Smadja & Butlin, 2009).

The only genera of Ithomiini known to sequester pyrrolizidine alkaloids as larvae from Apocynaceae host plants are *Tithorea* (Pliske, 1975; José R. Trigo, 2000; José Roberto Trigo & Brown, 1990). These compounds are usually gathered pharmacophagously from Asteraceae and Boraginaceae flowers by adult Ithomiini butterflies and are used for toxicity and pheromone production (Boppré, 1986; Edgar et al., 1976; Eisner & Meinwald, 1987; Stefan Schulz et al., 2004). Therefore, increased pheromone blend complexity in *Tithorea* may be associated with the ability to sequester plant compounds as larvae. Contrastingly, we do not find the same diversity of compounds in the closely-related species *Aeria eurimedia,* which also feeds on, but does not sequester compounds from, Apocynaceae larval host plants (José Roberto Trigo & Brown, 1990; Willmott & Freitas, 2006). Neither did we detect the same diversity of compounds in the androconial extracts of *Methona* species, which belong to the same early-diverging subtribe as *Tithorea* and *Aeria* (Tithoreina) and are suspected to sequester toxic compounds, from Solanaceae larval host plants (Elias et al., 2019; Massuda & Trigo, 2009). Increased *Tithorea* blend complexity in may therefore also be associated with the ability to sequester from Apocynaceae plants.

### Chemistry correlates with phylogeny

Chemical distances between species are significantly correlated with phylogenetic distances for both butterfly groups and tissue types. A similar correlation between chemical and genetic distances of geographically disparate *Heliconius* species has previously been found (Darragh et al., 2020). However, a similar comparison of variation in pyrrolizidine alkaloid derivatives across Ithomiini species found no phylogenetic signal (Stefan Schulz et al., 2004). The disconnect between chemistry and species relatedness in this latter study is attributed to the spatial and temporal variation in biotic conditions and pyrrolizidine alkaloid availability. It is therefore likely that we detected stronger phylogenetic signal in the current study because we collected our samples from the same locality during the same period. In addition, we incorporated all compounds that were consistently found within a species, including non-pyrrolizidine alkaloid derived compounds. Ithomiini male androconial extracts are dominated by pyrrolizidine alkaloid derivatives, and these are presumed to be the main signalling media (Ehlers & Schulz, 2022; Eisner & Meinwald, 1987; Stefan Schulz et al., 2004). However, recent studies have also found non-pyrrolizidine alkaloid related compounds in the androconial extracts of certain Ithomiini species (Mann et al., 2020; Stamm et al., 2019). The detection of phylogenetic signal in the current study would therefore suggest pheromone constituents are more diverse in this group than previously thought and the ability to produce pheromone compounds is also driven by shared history.

The observation of high phylogenetic signal suggests phenotypic similarity is the product of shared history, but does not allow for the distinction between evolutionary rates or processes, such as genetic drift, stabilising selection or punctuated divergent selection (Hansen & Martins, 1996; Revell et al., 2008). *Heliconius* androconial pheromones are involved in mate choice (Darragh et al., 2017) and have been shown to be most dissimilar between sister species with significant range overlap (Cama et al., 2022; González-Rojas et al., 2020). This suggests androconial pheromone divergence occurs via reproductive character displacement and facilitates reproductive isolation. Our results are consistent with these findings, given that high androconial chemical dissimilarity values are generally observed between sympatric species of low phylogenetic distance, whereas the same pattern is not observed among CSG extracts. If androconial compounds have been selected for species recognition, it is possible they have evolved via gradual shifts in pheromone composition, so that some aspects of the signal are conserved between closely related species (Buchinger & Li, 2020; Symonds & Elgar, 2008).

Divergence in *Heliconius* CSG chemistry has, instead, been suggested to be primarily driven by sexual selection for male-male competition (Catalina Estrada et al., 2011). Under a sexual selection regime, pressure for species recognition is relaxed. Therefore, while sexual selection may drive directional pheromone change in certain aspects of the signal (Steiger & Stökl, 2014), other features effective in the common ancestor are under no pressure to diverge and may be conserved. It is also possible that other evolutionary mechanisms contribute to chemical divergence. For example natural selection has reduced compound production in *Pieris* butterflies, to avoid chemical espionage by natural enemies (Huigens et al., 2010). Neutral evolution via genetic drift cannot be ruled out either, and it is possible some of the compounds analysed in this study are not involved in the signalling process, and therefore evolve primarily through neutral processes. Explicit tests of the function of different compounds and the fit of different evolutionary models will be required to understand the processes driving the evolution of heliconiine and ithomiine male butterfly chemistry.

## Conclusion

In this study we found chemical similarities between phylogenetically disparate species of butterflies that were not associated with mimicry. Overall, chemical profiles are highly species-specific and variation between species correlates with phylogenetic distance. This suggests chemical signalling is mostly used for intraspecific communication and chemical similarity between species is largely driven by shared ancestry. Patterns of occurrence in specific compounds shared between species suggest several selective pressures might be acting on male butterfly chemistry evolution. Further fitting of evolutionary models to butterfly pheromone data may shed more light on which selective processes are most influential. Furthermore, the field of avian olfaction is a growing one, and there is still much that is unknown. Additional research on the interaction between avian vision and olfaction will help us understand if and how chemical signals are used by toxic insects in predator avoidance.

## Author contributions

Rachel Blow, Stephanie Ehlers, Stefan Schulz, and Chris Jiggins were responsible for project conceptualization. Stephanie Ehlers, Daiane Szczerbowski, and Chris Jiggins collected samples. Rachel Blow and Chris Jiggins and designated species ring classifications. Rachel Blow, Stephanie Ehlers, Daiane Szczerbowski and Stefan Schulz conducted chemical analyses. Rachel Blow and Ian Warren conducted COI barcoding. Rachel Blow conducted statistical analyses and wrote the original manuscript. All authors contributed to manuscript revisions.

## Acknowledgements

We would like thank Keith Willmott for his invaluable help with butterfly species identification and guidance on mimicry ring classification. This study was supported by partnered funding from the National Environmental Research Council and Pembroke College, University of Cambridge. Additional thanks go to the Deutsche Forschungsgemeinschaft for providing funding. We are grateful to Joana Meier and Eva van der Heijden for providing species identifications for 29 ithomiine samples, prior to whole genome sequence deposition.

## Bibliography

1. Alkema, J. T., Dicke, M., & Wertheim, B. (2019). Context-Dependence and the Development of Push-Pull Approaches for Integrated Management of Drosophila suzukii. Insects, 10, 454. 10.3390/insects10120454

2. Altschul, S. F., Gish, W., Miller, W., Myers, E. W., & Lipman, D. J. (1990). Basic local alignment search tool. Journal of Molecular Biology, 215(3), 403–410. 10.1016/S0022-2836(05)80360-2

3. Avery, M., & Nelms, C. (1990). Food Avoidance by Red-Winged Blackbirds Conditioned with a Pyrazine Odor. The Auk, 107(3), 544–549. 10.1093/auk/107.3.544

4. Avilés, J. M., & Amo, L. (2018). The Evolution of Olfactory Capabilities in Wild Birds: A Comparative Study. Evolutionary Biology, 45(1), 27–36. 10.1007/s11692-017-9427-6

5. Bacquet, P. M. B., Brattström, O., Wang, H.-L., Allen, C. E., Löfstedt, C., Brakefield, P. M., & Nieberding, C. M. (2015). Selection on male sex pheromone composition contributes to butterfly reproductive isolation. Proceedings of the Royal Society B: Biological Sciences, 282, 20142734. 10.1098/rspb.2014.2734

6. Balthazart, J., & Taziaux, M. (2009). The underestimated role of olfaction in avian reproduction? Behavioural Brain Research, 200(2), 248–259. 10.1016/j.bbr.2008.08.036

7. Beccaloni, G. W. (1997). Ecology, Natural History and Behaviour of Ithomiine Butterflies and Their Mimics in Ecuador (Lepidoptera: Nymphalidae: Ithomiinae). Tropical Lepidoptera., 8(2), 103–124.

8. Benson, D. A., Cavanaugh, M., Clark, K., Karsch-Mizrachi, I., Lipman, D. J., Ostell, J., & Sayers, E. W. (2013). GenBank. Nucleic Acids Research, 41(D1). 10.1093/nar/gks1195

9. Birch, M. C., & Haynes, K. F. (1982). Insect pheromones. In edward Arnold. Edward Arnold. 10.2307/2403157

10. Bonacci, T., Brandmayr, P., Dalpozzo, R., De Nino, A., Massolo, A., Tagarelli, A., & Brandmayr, T. Z. (2008). Odour and colour similarity in two species of gregarious carabid beetles (Coleoptera) from the Crati Valley, southern Italy: A case of Müllerian mimicry? Entomological News, 119(4), 325–337. 10.3157/0013-872X-119.4.325

11. Boppré, M. (1986). Insects pharmacophagously utilizing defensive plant chemicals (pyrrolizidine alkaloids). Natur Wissenscahften, 26, 17–26.

12. Bosque, R. J., Hyseni, C., Santos, M. L. G., Rangel, E., Da Silva Dias, C. J., Hearin, J. B., Da Silva, N. J., Domingos, F. M. C. B., Colli, G. R., & Noonan, B. P. (2022). Müllerian mimicry and the coloration patterns of sympatric coral snakes. Biological Journal of the Linnean Society, 135(4), 645–651. 10.1093/biolinnean/blab155

13. Brower, L. P., Brower, J. V. Z., & Collins, C. T. (1963). Experimental studies of mimicry. 7. Relative palatability and Müllerian mimicry among neotropical butterflies of the subfamily Heliconiinae. In Zoologica : scientific contributions of the New York Zoological Society. (Vol. 48, Issue 7, pp. 65–84). New York,. 10.5962/p.203311

14. Buchinger, T. J., & Li, W. (2020). The evolution of (non)species-specific pheromones. Evolutionary Ecology, 34(4), 455–468. 10.1007/s10682-020-10046-0

15. Bura, V. L., Fleming, A. J., & Yack, J. E. (2009). What’s the buzz? Ultrasonic and sonic warning signals in caterpillars of the great peacock moth (Saturnia pyri). Naturwissenschaften, 96(6), 713–718. 10.1007/s00114-009-0527-8

16. Burdfield-Steel, E., Pakkanen, H., Rojas, B., Galarza, J. A., & Mappes, J. (2018). De novo synthesis of chemical defenses in an aposematic moth. Journal of Insect Science, 18(2). 10.1093/jisesa/iey020

17. Byers, K. J. R. P., Darragh, K., Musgrove, J., Almeida, D. A., Garza, S. F., Warren, I. A., Rastas, P. M., Kučka, M., Chan, Y. F., Merrill, R. M., Schulz, S., McMillan, W. O., & Jiggins, C. D. (2020). A major locus controls a biologically active pheromone component in Heliconius melpomene. Evolution, 74(2), 349–364. 10.1111/evo.13922

18. Calcagnile, M., Tredici, S. M., Talà, A., & Alifano, P. (2019). Bacterial Semiochemicals and Transkingdom Interactions with Insects and Plants. Insects, 10. 10.3390/insects10120441

19. Cama, B., Ehlers, S., Szczerbowski, D., Thomas-Oates, J., Jiggins, C. D., Schulz, S., McMillan, W. O., & Dasmahapatra, K. K. (2022). Exploitation of an ancestral pheromone biosynthetic pathway contributes to diversification in Heliconius butterflies. Proceedings of the Royal Society B: Biological Sciences, 289(1979). 10.1098/rspb.2022.0474

20. Chazot, N., Willmott, K. R., Lamas, G., Freitas, A. V. L., Piron-Prunier, F., Arias, C. F., Mallet, J., De-Silva, D. L., & Elias, M. (2019). Renewed diversification following Miocene landscape turnover in a Neotropical butterfly radiation. Global Ecology and Biogeography, 28(8), 1118–1132. 10.1111/geb.12919

21. Cicconardi, F., Milanetti, E., Pinheiro de Castro, E. C., Mazo-Vargas, A., Van Belleghem, S. M., Ruggieri, A. A., Rastas, P., Hanly, J., Evans, E., Jiggins, C. D., Owen McMillan, W., Papa, R., Di Marino, D., Martin, A., & Montgomery, S. H. (2023). Evolutionary dynamics of genome size and content during the adaptive radiation of Heliconiini butterflies. Nature Communications, 14(1). 10.1038/s41467-023-41412-5

22. Conner, W. E., & Iyengar, V. K. (2016). Male Pheromones in Moths: Reproductive Isolation, Sexy Sons, and Good Genes. In Pheromone Communication in Moths (pp. 191–208). 10.1525/9780520964433-013

23. Crane, J. (1955). Imaginal behaviour of a Trinidad butterfly, Heliconius erato hydara Hewitson, with special reference to the social use of color. Zoologica, II(1–18), 167–194.

24. Darragh, K., Jiggins, C. D., Vanjari, S., Mann, F., Gonzalez-Rojas, M. F., Morrison, C. R., Salazar, C., Pardo-Diaz, C., Merrill, R. M., Mcmillan, W. O., & Schulz, S. (2017). Male sex pheromone components in Heliconius butterflies released by the androconia affect female choice. PeerJ, e3953. 10.7717/peerj.3953

25. Darragh, K., Montejo-Kovacevich, G., Kozak, K. M., Morrison, C. R., Figueiredo, C. M. E., Ready, J. S., Salazar, C., Linares, M., Byers, K. J. R. P., Merrill, R. M., McMillan, W. O., Schulz, S., & Jiggins, C. D. (2020). Species specificity and intraspecific variation in the chemical profiles of Heliconius butterflies across a large geographic range. Ecology and Evolution, 10(9), 3895–3918. 10.1002/ece3.6079

26. Dettner, K., & Liepert, C. (1994). Chemical mimicry and camouflage. Annual Review of Entomology, 39(c), 129–154. 10.1146/annurev.en.39.010194.001021

27. Edgar, J., Culvenor, C., & Pliske, T. (1976). Isolation of a lactone, structurally related to the esterifying acids of pyrrolizdine alkaloids, from the costal fringes of male Ithomiinae. Journal of Chemical Ecology, 2(Ii), 263–270.

28. Ehlers, S., & Schulz, S. (2022). The scent chemistry of butterflies. In Natural Product Reports (Vol. 40, Issue 4, pp. 794–818). 10.1039/d2np00067a

29. Ehlers, S., Szczerbowski, D., Harig, T., Stell, M., Hötling, S., Darragh, K., Jiggins, C. D., & Schulz, S. (2021). Identification and Composition of Clasper Scent Gland Components of the Butterfly Heliconius erato and Its Relation to Mimicry. ChemBioChem, 22, 3300– 3313. 10.1002/cbic.202100372

30. Eisner, T. (1970). Chemical Defense against Predation in Arthropods. In E. Sondheimer & J. Simeone (Eds.), Chemical Ecology (pp. 157–217). Academic Press.

31. Eisner, T., & Meinwald, J. (1987). Alkaloid-Derived Pheromones and Sexual Selection in Lepidoptera. In Pheromone Biochemistry (pp. 251–269). ACADEMIC PRESS, INC. 10.1016/b978-0-12-564485-3.50013-0

32. Eisner, T., & Meinwald, J. (1995). The chemistry of sexual selection. Proceedings of the National Academy of Sciences of the United States of America, 92(1), 50–55. 10.1073/pnas.92.1.50

33. El-Sayed, A. M. (2023). The Pherobase: Database of Pheromones and Semiochemicals. <http://Www.Pherobase.Com>.

34. Elias, M., Gompert, Z., Jiggins, C., & Willmott, K. (2008). Mutualistic interactions drive ecological niche convergence in a diverse butterfly community. PLoS Biology, 6(12). 10.1371/journal.pbio.0060300

35. Elias, M., Mcclure, M., Clerc, C., Desbois, C., Meichanetzoglou, A., Cau, M., Bastin-Héline, L., Bacigalupo, J., Houssin, C., Pinna, C., Nay, B., Llaurens, V., Berthier, S., Andraud, C., & Gomez, D. (2019). Why has transparency evolved in aposematic butterflies? Insights from the largest radiation of aposematic butterflies, the Ithomiini. Proceedings of the Royal Society B: Biological Sciences, 286, 20182769. 10.1098/rspb.2018.2769

36. Estrada, C., & Jiggins, C. D. (2008). Interspecific sexual attraction because of convergence in warning colouration: Is there a conflict between natural and sexual selection in mimetic species? Journal of Evolutionary Biology, 21(3), 749–760. 10.1111/j.1420-9101.2008.01517.x

37. Estrada, Catalina, Schulz, S., Yildizhan, S., & Gilbert, L. E. (2011). Sexual selection drives the evolution of antiaphrodisiac pheromones in butterflies. Evolution, 65(10), 2843– 2854.

38. Estrada, Catalina, Yildizhan, S., Schulz, S., & Gilbert, L. E. (2010). Sex-specific chemical cues from immatures facilitate the evolution of mate guarding in Heliconius butterflies. Proceedings of the Royal Society B: Biological Sciences, 277(1680), 407–413. 10.1098/rspb.2009.1476

39. Funaro, C. F., Böröczky, K., Vargo, E. L., & Schal, C. (2018). Identification of a queen and king recognition pheromone in the subterranean termite Reticulitermes flavipes. Proceedings of the National Academy of Sciences of the United States of America, 115(15), 3888–3893. 10.1073/pnas.1721419115

40. Gilbert, L. E. (1976). Postmating Female Odor in Heliconius Butterflies: A Male-Contributed Antiaphrodisiac? Science, 193(4251), 419–420.

41. Gomez-Diaz, C., & Benton, R. (2013). The joy of sex pheromones. EMBO Reports, 14(10), 874–883. 10.1038/embor.2013.140

42. González-Rojas, M. F., Darragh, K., Robles, J., Linares, M., Schulz, S., McMillan, W. O., Jiggins, C. D., Pardo-Diaz, C., & Salazar, C. (2020). Chemical signals act as the main reproductive barrier between sister and mimetic Heliconius butterflies. Proceedings of the Royal Society B: Biological Sciences, 287(1926), 20200587. 10.1098/rspb.2020.0587

43. Grieves, L. A., Gilles, M., Cuthill, I. C., Székely, T., MacDougall-Shackleton, E. A., & Caspers, B. A. (2022). Olfactory camouflage and communication in birds. Biological Reviews, 97(3), 1193–1209. 10.1111/brv.12837

44. Groot, A. T., Dekker, T., & Heckel, D. G. (2016). The Genetic Basis of Pheromone Evolution in Moths. Annual Review of Entomology, 61(1), 99–117. 10.1146/annurev-ento-010715-023638

45. Groot, A. T., Horovitz, J. L., Hamilton, J., Santangelo, R. G., Schal, C., & Gould, F. (2006). Experimental evidence for interspecific directional selection on moth pheromone communication. www.pnas.orgcgidoi10.1073pnas.0508609103

46. Guilford, T., Nicol, C., Rothschild, M., & Moore, B. P. (1987). The biological roles of pyrazines: evidence for a warning odour function. Biological Journal of the Linnean Society, 31(2), 113–128. 10.1111/j.1095-8312.1987.tb01984.x

47. Hansen, T. F., & Martins, E. P. (1996). Translating Between Microevolutionary Process and Macroevolutionary Patterns: The Correlation Structure of Interspecific Data. In Source: Evolution (Vol. 50, Issue 4).

48. Hebets, E. A., & Papaj, D. R. (2005). Complex signal function: Developing a framework of testable hypotheses. Behavioral Ecology and Sociobiology, 57(3), 197–214. 10.1007/s00265-004-0865-7

49. Hill, R. I. (2021). Convergent flight morphology among Müllerian mimic mutualists. Evolution, 75, 2460–2479. 10.1111/evo.14331

50. Honda, K. (1981). Larval osmeterial secretions of the swallowtails (Papilio). Journal of Chemical Ecology, 7(6), 1089–1113. 10.1007/BF00987631

51. Huigens, M. E., Woelke, J. B., Pashalidou, F. G., Bukovinszky, T., Smid, H. M., & Fatouros, N. E. (2010). Chemical espionage on species-specific butterfly anti-aphrodisiacs by hitchhiking Trichogramma wasps. Behavioral Ecology, 21(3), 470–478. 10.1093/beheco/arq007

52. Jiggins, C. D. (2017). The Ecology and Evolution of Heliconius Butterflies. Oxford University Press.

53. Jiggins, C. D., Estrada, C., & Rodrigues, A. (2004). Mimicry and the evolution of premating isolation in Heliconlus melpomene Linnaeus. Journal of Evolutionary Biology, 17(3), 680–691. 10.1111/j.1420-9101.2004.00675.x

54. Kaae, R., Shorey, H. H., & Gaston, L. K. (1973). Pheromone Concentration as a Mechanism for Reproductive Isolation between Two Lepidopterous Species. Science, 179, 487–488. 10.1126/science.179.4072.487

55. Kozak, K. M., Wahlberg, N., Neild, A. F. E., Dasmahapatra, K. K., Mallet, J., & Jiggins, C. D. (2015). Multilocus species trees show the recent adaptive radiation of the mimetic heliconius butterflies. Systematic Biology, 64(3), 505–524. 10.1093/sysbio/syv007

56. Langham, G. M. (2004). Specialized avian predators repeatedly attack novel color morphs of Heliconius butterflies. In Evolution (Vol. 58, Issue 12).

57. Langham, G. M. (2006). Rufous-tailed jacamars and aposematic butterflies: Do older birds attack novel prey? Behavioral Ecology, 17(2), 285–290. 10.1093/beheco/arj027

58. Leoncini, I., Le Conte, Y., Costagliola, G., Plettner, E., Toth, A. L., Wang, M., Huang, Z., Bécard, J. M., Crauser, D., Slessor, K. N., & Robinson, G. E. (2004). Regulation of behavioral maturation by a primer pheromone produced by adult worker honey bees. Proceedings of the National Academy of Sciences of the United States of America, 101(50), 17559–17564. 10.1073/pnas.0407652101

59. Löfstedt, C., Herrebout, W. M., & Menken, S. B. J. (1991). Sex pheromones and their potential role in the evolution of reproductive isolation in small ermine moths (Yponomeutidae). Chemoecology, 2(1), 20–28. 10.1007/BF01240662

60. Mallard, G., & Reed, J. (1997). Automated Mass Spectral Deconvolution & Identification System. AMDIS User Guide. 10.1159/000249215

61. Mallet, J., Barton, N., Santisteban, J. C., Muedas, M. M., & Eeley, H. (1990). Estimates of Selection and Gene Flow From Measures of Cline Width and Linkage Disequilibrium in Heliconius Hybrid Zones. https://academic.oup.com/genetics/article/124/4/921/5999976

62. Mallet, J., & Gilbert, L. E. (1995). Why are there so many mimicry rings? Correlations between habitat, behaviour and mimicry in Heliconius butterflies. Biological Journal of the Linnean Society, 55(2), 159–180. 10.1111/j.1095-8312.1995.tb01057.x

63. Malouines, C. (2017). Counter-perfume: using pheromones to prevent female remating. Biological Reviews, 92(3), 1570–1581. 10.1111/brv.12296

64. Manger, P., Corfield, J. R., Price, K., Iwaniuk, A. N., Gutiérrez-Ibáñez, C., Birkhead, T., & Wylie, D. R. (2015). Diversity in olfactory bulb size in birds reflects allometry, ecology, and phylogeny. Frontiers in Neuroanatomy, 9, 102. 10.3389/fnana.2015.00102

65. Mann, F., Szczerbowski, D., De Silva, L., Mcclure, M., Elias, M., & Schulz, S. (2020). 3-Acetoxy-fatty acid isoprenyl esters from androconia of the ithomiine butterfly Ithomia salapia. Beilstein J. Org. Chem, 16, 2776–2787. 10.3762/bjoc.16.228

66. Mann, F., Vanjari, S., Rosser, N., Mann, S., Dasmahapatra, K. K., Corbin, C., Linares, M., Pardo-Diaz, C., Salazar, C., Jiggins, C., & Schulz, S. (2017). The Scent Chemistry of Heliconius Wing Androconia. Journal of Chemical Ecology, 43(9), 843–857. 10.1007/s10886-017-0867-3

67. Mappes, J., Marples, N., & Endler, J. A. (2005). The complex business of survival by aposematism. Trends in Ecology and Evolution, 20(11), 598–603. 10.1016/j.tree.2005.07.011

68. Massuda, K. F., & Trigo, J. R. (2009). Chemical defence of the warningly coloured caterpillars of Methona themisto (Lepidoptera: Nymphalidae: Ithomiinae). European Journal of Entomology, 106(2), 253–259. 10.14411/eje.2009.033

69. Melo, D. J., Borges, E. O., Szczerbowski, D., Vidal, D. M., Schulz, S., & Zarbin, P. H. G. (2022). Identification and Synthesis of a Macrolide as an Anti-aphrodisiac Pheromone from Males of Heliconius erato phyllis. Organic Letters, 24(21), 3772–3775. 10.1021/acs.orglett.2c01160

70. Mérot, C., Frérot, B., Leppik, E., & Joron, M. (2015). Beyond magic traits: Multimodal mating cues in Heliconius butterflies. Source: Evolution, 69(11), 2891–2904. 10.1111/evo

71. Mérot, C., Mavárez, J., Evin, A., Dasmahapatra, K. K., Mallet, J., Lamas, G., & Joron, M. (2013). Genetic differentiation without mimicry shift in a pair of hybridizing Heliconius species (Lepidoptera: Nymphalidae). In Biological Journal of the Linnean Society. 10.1111/bij.12091

72. Mitaka, Y., & Akino, T. (2021). A Review of Termite Pheromones: Multifaceted, Context-Dependent, and Rational Chemical Communications. Frontiers in Ecology and Evolution, 8, 595614. 10.3389/fevo.2020.595614

73. Moore, B. P., Brown, W. V., & Rothschild, M. (1990). Methylalkylpyrazines in aposematic insects, their hostplants and mimics. In Chemoecology (Vol. 1, Issue 2, pp. 43–51). 10.1007/BF01325227

74. Morgan, M., Anders, S., Lawrence, M., Aboyoun, P., Pagès, H., & Gentleman, R. (2009). “ShortRead: a Bioconductor package for input, quality assessment and exploration of high-throughput sequence data.” Bioinformatics, 25, 2607–2608. 10.1093/bioinformatics/btp450

75. Nago, H., & Matsumoto, M. (1994). An Ecological Role of Volatiles Produced by Lasiodiplodia theobromae. Bioscience, Biotechnology, and Biochemistry, 58(7), 1267– 1272. 10.1271/bbb.58.1267

76. Nakashima, Y., Birkett, M. A., Pye, B. J., & Powell, W. (2006). Chemically mediated intraguild predator avoidance by aphid parasitoids: Interspecific variability in sensitivity to semiochemical trails of ladybird predators. Journal of Chemical Ecology, 32(9), 1989–1998. 10.1007/s10886-006-9123-y

77. Nishida, R., Schulz, S., Kim, C. S., Fukami, H., Kuwahara, Y., Honda, K., & Hayashi, N. (1996). Male sex pheromone of a giant danaine butterfly, Idea leuconoe. Journal of Chemical Ecology, 22(5), 949–972. 10.1007/BF02029947

78. O’Reilly, L. J., Agassiz, D. J. L., Neil, T. R., & Holderied, M. W. (2019). Deaf moths employ acoustic Müllerian mimicry against bats using wingbeat-powered tymbals. Scientific Reports, 9(1), 1–9. 10.1038/s41598-018-37812-z

79. Oksanen, J., Simpson, G., Blanchet, F., Kindt, R., Legendre, P., Minchin, P., O’Hara, R., Solymos, P., Stevens, M., Szoecs, E., Wagner, H., Barbour, M., Bedward, M., Bolker, B., Borcard, D., Carvalho, G., Chirico, M. M. D. C., Durand, S., … Weedon, J. (2022). vegan: Community Ecology Package. R Package Version, 2.6–4. https://cran.r-project.org/package=vegan

80. Orlova, M., & Amsalem, E. (2021). Bumble bee queen pheromones are context-dependent. Scientific Reports, 11(1), 1–7. 10.1038/s41598-021-96411-7

81. Palma-Onetto, V., Bergmann, J., & González-Teuber, M. (2023). Mode of action, chemistry and defensive efficacy of the osmeterium in the caterpillar Battus polydamas archidamas. Scientific Reports |, 13, 6644. 10.1038/s41598-023-33390-x

82. Paradis, E., & Schliep, K. (2019). ape 5.0: an environment for modern phylogenetics and evolutionary analyses in R. Bioinformatics, 35, 526–528. 10.1093/bioinformatics/bty633

83. Pasteels, J. M., Gregoire, J.-C., & Rowell-Rahier, M. (1983). The chemical ecology of defense in arthropods. Ann. Rev. EntomoL, 28, 263–289. www.annualreviews.org

84. Pinheiro, C. E. G. (1996). Palatablility and escaping ability in N eotropical butterflies: tests with wild kingbirds (Tyrannus rnelancholicus, Tyrannidae). https://academic.oup.com/biolinnean/article/59/4/351/2705811

85. Pinheiro, C. E. G. (2003). Does Müllerian Mimicry Work in Nature? Experiments with Butterflies and Birds (Tyrannidae). Biotropica, 35(3), 356–364. 10.1111/j.1744-7429.2003.tb00589.x

86. Pinheiro, C. E. G., Medri, Í. M., & Salcedo, A. K. M. (2008). Why do the ithomiines (Lepidoptera, Nymphalidae) aggregate? Notes on a butterfly pocket in central Brazil. Revista Brasileira de Entomologia, 52(4), 610–614. 10.1590/S0085-56262008000400012

87. Pliske, T. (1975). Courthsip behaviour and use of chemical communication by males of certain species of Ithomiine butterflies (Nymphalidae:Lepidoptera). Annals of the Entomological Society of America, 935–942.

88. R Core Team. (2023). R: A Language and Environment for Statistical Computing. https://www.r-project.org/

89. Revell, L. J., Harmon, L. J., & Collar, D. C. (2008). Phylogenetic Signal, Evolutionary Process, and Rate. Syst. Biol, 57(4), 591–601. 10.1080/10635150802302427

90. Rothschild, M. (1961). Defensive odours and Mullerian mimicry among insects. 113, 101– 113.

91. Rowe, C., & Guilford, T. (1999). The evolution of multimodal warning displays. Evolutionary Ecology, 13(7–8), 655–671. 10.1023/A:1011021630244

92. Rowe, C., & Halpin, C. (2013). Why are warning displays multimodal? Behavioral Ecology and Sociobiology, 67(9), 1425–1439. 10.1007/s00265-013-1515-8

93. Rowe, C., & Skelhorn, J. (2004). Avian psychology and communication. Proceedings of the Royal Society B: Biological Sciences, 271, 1435–1442. 10.1098/rspb.2004.2753

94. Rubene, D., Low, M., & Brodin, A. (2023). Birds differentially prioritize visual and olfactory foraging cues depending on habitat of origin and sex. 10.1098/rsos.221336

95. Ruxton, G. D., Allen, W. L., Sherratt, T. N., & Speed, M. P. (2018). Avoiding Attack: the evolutionary ecology of crypsis, aposematism, and mimicry, 2nd edn (Vol. 1). Oxford University Press. 10.1093/oso/9780199688678.001.0001

96. Salcedo, C. (2011). Evidence of predation and disturbance events at Heliconius (Insecta: Lepidoptera: Nymphalidae) nocturnal aggregations in Panama and Costa Rica. Journal of Natural History, 45, 1715–1721. 10.1080/00222933.2011.559692

97. Schulz, S, Francke, W., Edgar, J., & Schneider, D. (1988). Volatile Compounds from Androconial Organs of Danaine and Ithomiine Butterflies. In Z. Naturforsch (Vol. 43).

98. Schulz, Stefan. (2009). Alkaloid-derived male courtship pheromones. In W. E. Conner (Ed.), Tiger moths and woolly bears : behavior, ecology, and evolution of the Arctiidae (pp. 145–153).

99. Schulz, Stefan, Beccaloni, G., Brown, K. S., Boppré, M., Freitas, A. V. L., Ockenfels, P., & Trigo, J. R. (2004). Semiochemicals derived from pyrrolizidine alkaloids in male ithomiine butterflies (Lepidoptera: Nymphalidae: Ithomiinae). Biochemical Systematics and Ecology, 32(8), 699–713. 10.1016/j.bse.2003.12.004

100. Schulz, Stefan, Estrada, C., Yildizhan, S., Boppré, M., & Gilbert, L. E. (2008). An Antiaphrodisiac in Heliconius melpomene Butterflies. Journal of Chemical Ecology, 34, 82–93. 10.1007/s10886-007-9393-z

101. Shahandeh, M. P., Pischedda, A., & Turner, T. L. (2018). Male mate choice via cuticular hydrocarbon pheromones drives reproductive isolation between Drosophila species. Evolution, 72(1), 123–135. 10.1111/evo.13389

102. Sherratt, T. N. (2008). The evolution of Mullerian Mimicry. In Nature (Vol. 388, pp. 539– 547).

103. Smadja, C., & Butlin, R. K. (2009). On the scent of speciation: The chemosensory system and its role in premating isolation. In Heredity (Vol. 102, Issue 1, pp. 77–97). 10.1038/hdy.2008.55

104. Srygley, R. B. (1999). Locomotor mimicry in Heliconius butterflies: Contrast analyses of flight morphology and kinematics. Philosophical Transactions of the Royal Society B: Biological Sciences, 354(1380), 203–214. 10.1098/rstb.1999.0372

105. Stamm, P., Mann, F., Mcclure, M., Elias, M., & Schulz, S. (2019). Chemistry of the Androconial Secretion of the Ithomiine Butterfly Oleria onega. Journal of Chemical Ecology, 45, 768–778. 10.1007/s10886-019-01100-5

106. Steiger, S. S., Fidler, A. E., Valcu, M., & Kempenaers, B. (2008). Avian olfactory receptor gene repertoires: Evidence for a well-developed sense of smell in birds? Proceedings of the Royal Society B: Biological Sciences, 275(1649), 2309–2317. 10.1098/rspb.2008.0607

107. Steiger, S., & Stökl, J. (2014). The Role of Sexual Selection in the Evolution of Chemical Signals in Insects. Insects, 5(2), 423–438. 10.3390/insects5020423

108. Sugiura, S. (2020). Predators as drivers of insect defenses. Entomological Science, 23, 316–337.

109. Symonds, M. R. E., & Elgar, M. A. (2008). The evolution of pheromone diversity. Trends in Ecology and Evolution, 23(4), 220–228. 10.1016/j.tree.2007.11.009

110. Symonds, M. R. E., Johnson, T. L., & Elgar, M. A. (2011). Pheromone production, male abundance, body size, and the evolution of elaborate antennae in moths. Ecology and Evolution, 2(1), 227–246. 10.1002/ece3.81

111. Symula, R., Schulte, R., & Summers, K. (2001). Molecular phylogenetic evidence for a mimetic radiation in Peruvian poison frogs supports a Müllerian mimicry hypothesis. Proceedings of the Royal Society B: Biological Sciences, 268(1484), 2415–2421. 10.1098/rspb.2001.1812

112. Trigo, José R. (2000). The Chemistry of Antipredator Defense by Secondary Compounds in Neotropical Lepidoptera: Facts, Perspectives and Caveats. Journal of the Brazilian Chemical Society, 11(6), 551–561. 10.1590/S0103-50532000000600002

113. Trigo, José Roberto, & Brown, K. S. (1990). Variation of pyrrolizidine alkaloids in Ithomiinae: a comparative study between species feeding on Apocynaceae and Solanaceae. In Chemoecology (Vol. 22).

114. Vencl, F. V, Ottens, K., Dixon, M. M., Candler, S., Bernal, X. E., Estrada, C., & Page, R. A. (2016). Pyrazine emission by a tropical firefly. 48(5), 645–655. 10.2307/48577089

115. Wang, L.-M., Li, N., Zhang, M., Tang, Q., Lu, H.-Z., Zhou, Q.-Y., Niu, J.-X., Xiao, L., Peng, Z.-Y., Zhang, C., Liu, M., Wang, D.-Q., & Deng, S.-Q. (2023). The sex pheromone heptacosane enhances the mating competitiveness of sterile Aedes aegypti males. Parasites & Vectors, 16, 1–9. 10.1186/s13071-023-05711-6

116. Wicker-Thomas, C. (2011). Evolution of insect pheromones and their role in reproductive isolation and speciation. Annales de La Societe Entomologique de France, 47(1–2), 55–62. 10.1080/00379271.2011.10697696

117. Wickham, H. (2016). ggplot2: Elegant Graphics for Data Analysis. Springer-Verlag New York. https://ggplot2.tidyverse.org

118. Willmott, K. R., & Freitas, A. V. L. (2006). Higher-level phylogeny of the Ithomiinae (Lepidoptera: Nymphalidae): Classification, patterns of larval hostplant colonization and diversification. Cladistics, 22(4), 297–368. 10.1111/j.1096-0031.2006.00108.x

119. Willmott, K. R., & Mallet, J. (2004). Correlations between adult mimicry and larval host plants in ithomiine butterflies. Proceedings of the Royal Society B: Biological Sciences, 271(SUPPL. 5). 10.1098/rsbl.2004.0184

120. Wyatt, T. D. (2014). Pheromones and Animal behaviour: chemical signals and signatures. Cambridge University Press.

121. Yang, C. Y., Han, K. S., & Boo, K. S. (2009). Sex pheromones and reproductive isolation of three species in genus Adoxophyes. Journal of Chemical Ecology, 35(3), 342–348. 10.1007/s10886-009-9602-z

122. Yang, S., Zhang, X. F., Gao, Y. L., Chen, D., She, D. M., Zhang, T., & Ning, J. (2017). Male-Produced Aggregation Pheromone of Coffee Bean Weevil, Araecerus fasciculatus. Journal of Chemical Ecology, 43(10), 978–985. 10.1007/s10886-017-0894-0

123. Yew, J. Y., & Chung, H. (2015). Insect pheromones: An overview of function, form, and discovery. In Progress in Lipid Research (Vol. 59, pp. 88–105). 10.1016/j.plipres.2015.06.001

124. Zhang, J. P., Salcedo, C., Fang, Y. L., Zhang, R. J., & Zhang, Z. N. (2012). An overlooked component: (Z)-9-tetradecenal as a sex pheromone in Helicoverpa armigera. Journal of Insect Physiology, 58(9), 1209–1216. 10.1016/j.jinsphys.2012.05.018

