## Supplementary Material for "Phylogeny not mimicry drives chemical variation in tropical aposematic butterflies"

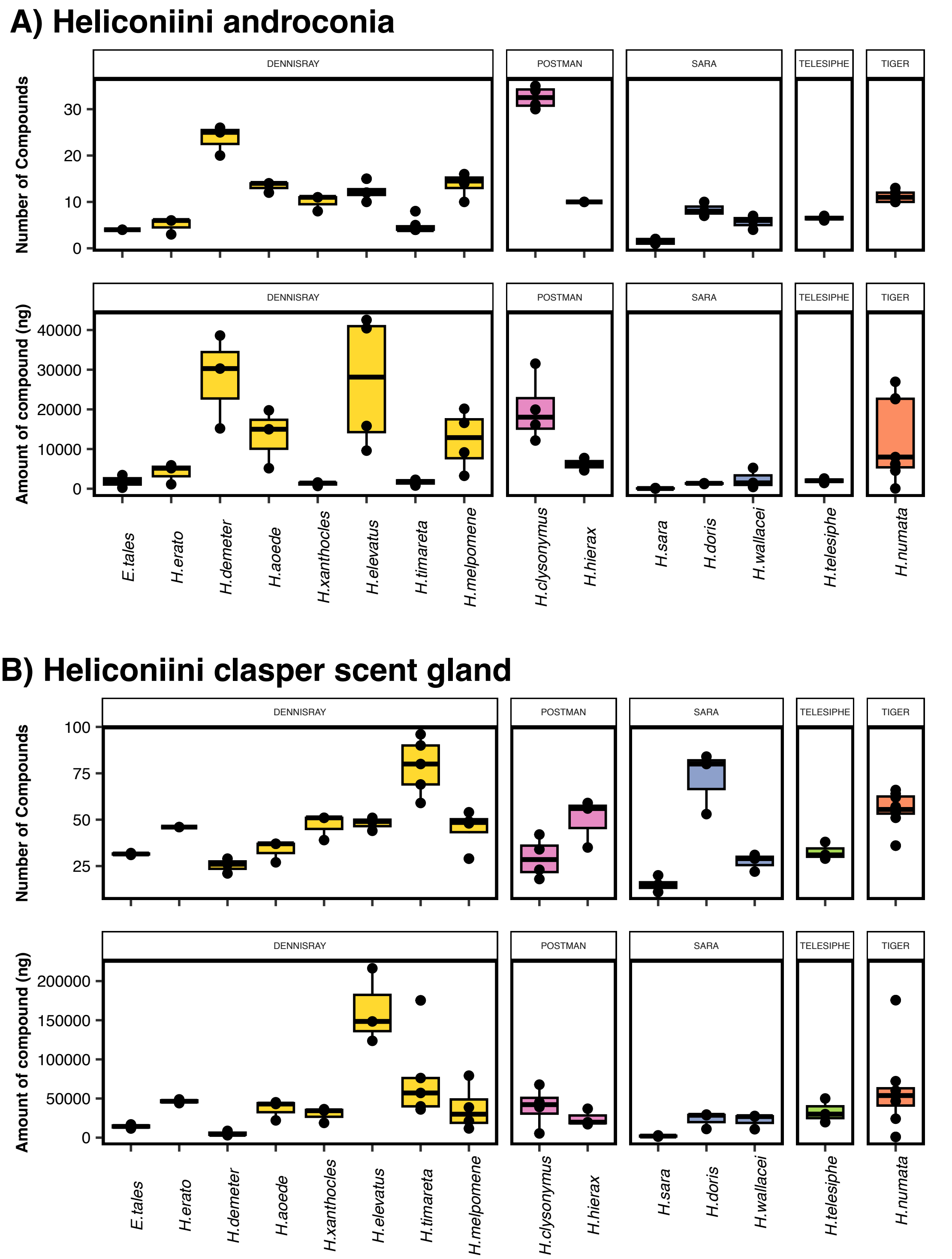


**Figure S1.** Boxplot displaying variation in the number and total amount (nanograms) of Heliconiini androconial (A) and clasper scent gland (B) compounds between species. Boxes are coloured and separated according to mimicry, and points represent individual measurements within species.

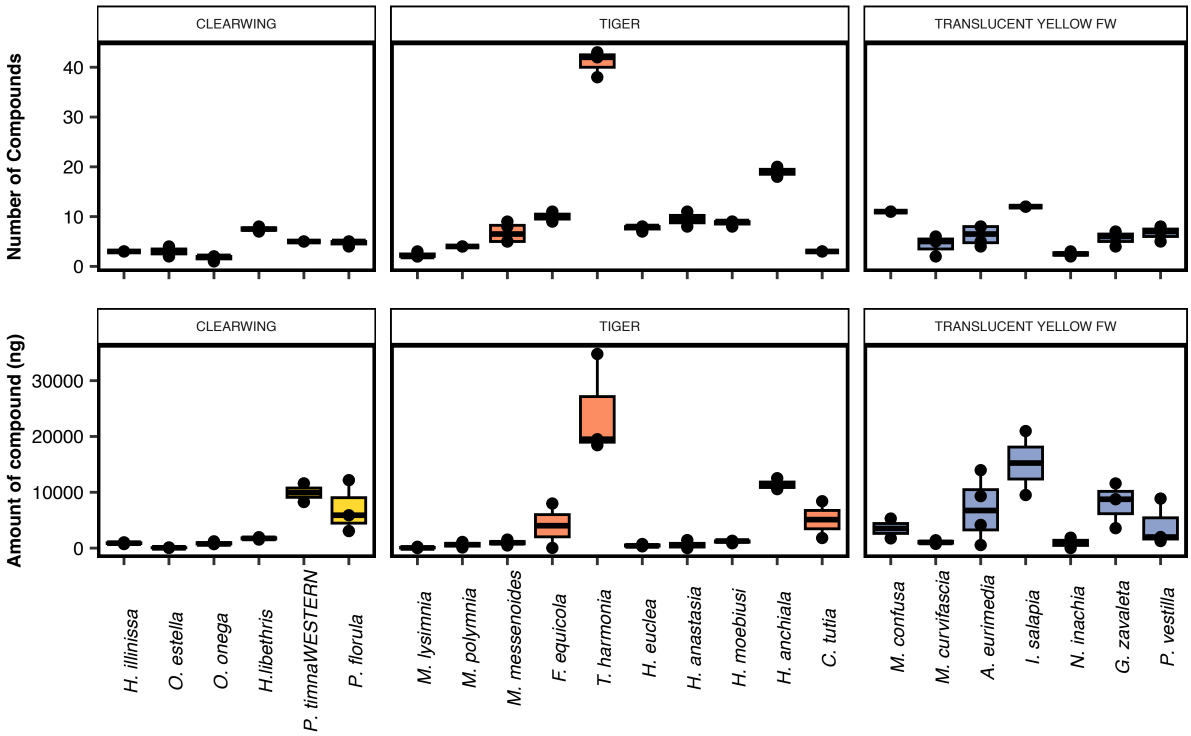


**Figure S2.** Boxplot displaying variation in the number and total amount (in nanograms) of Ithomiini androconial compounds between species. Boxes are coloured and separated according to mimicry and points represent individual measurements within species.

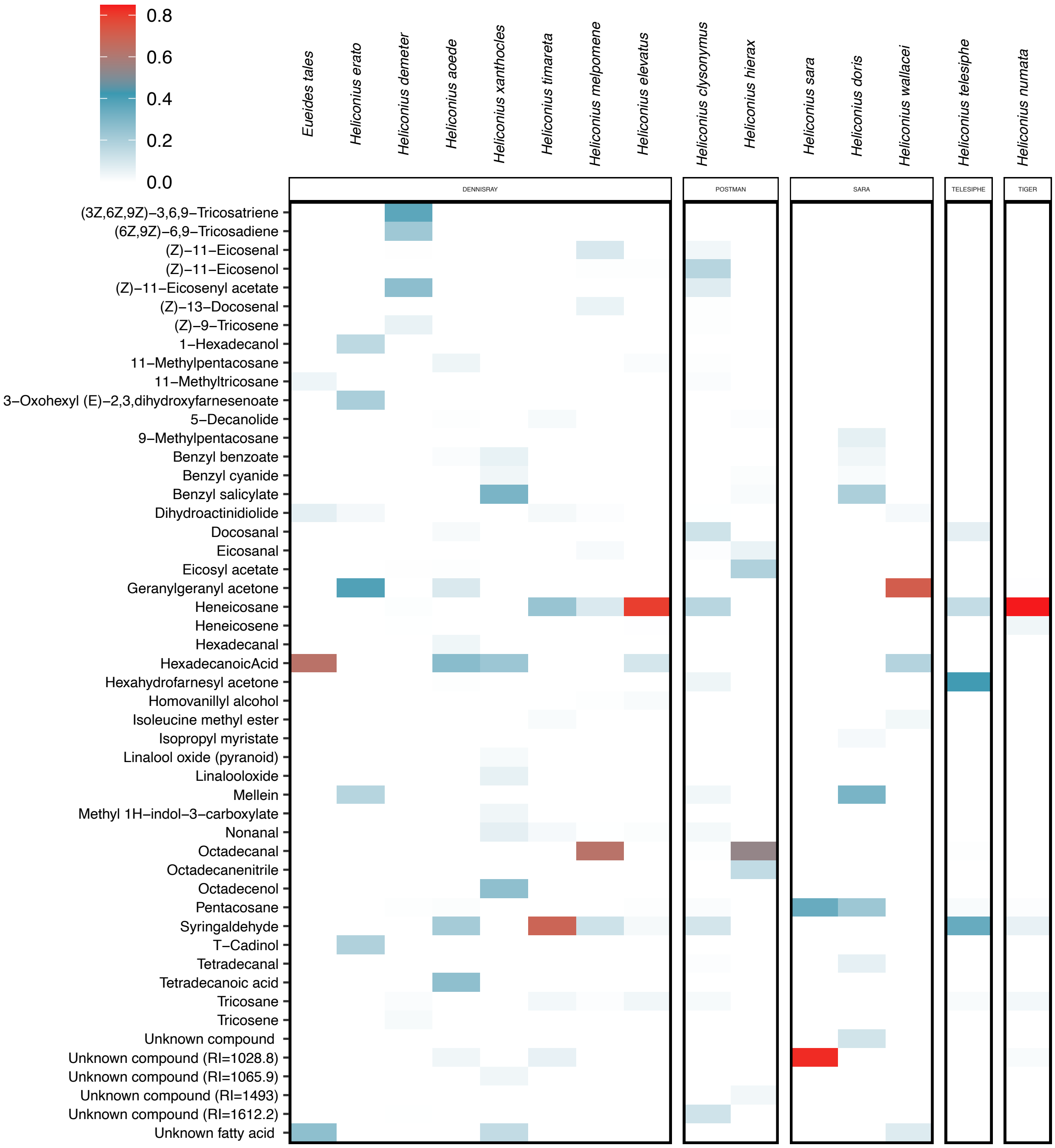


**Figure S3.** Variation in absolute compound concentration (nanograms) among the 50 most abundant Heliconiini androconial compounds, with species grouped by mimicry.

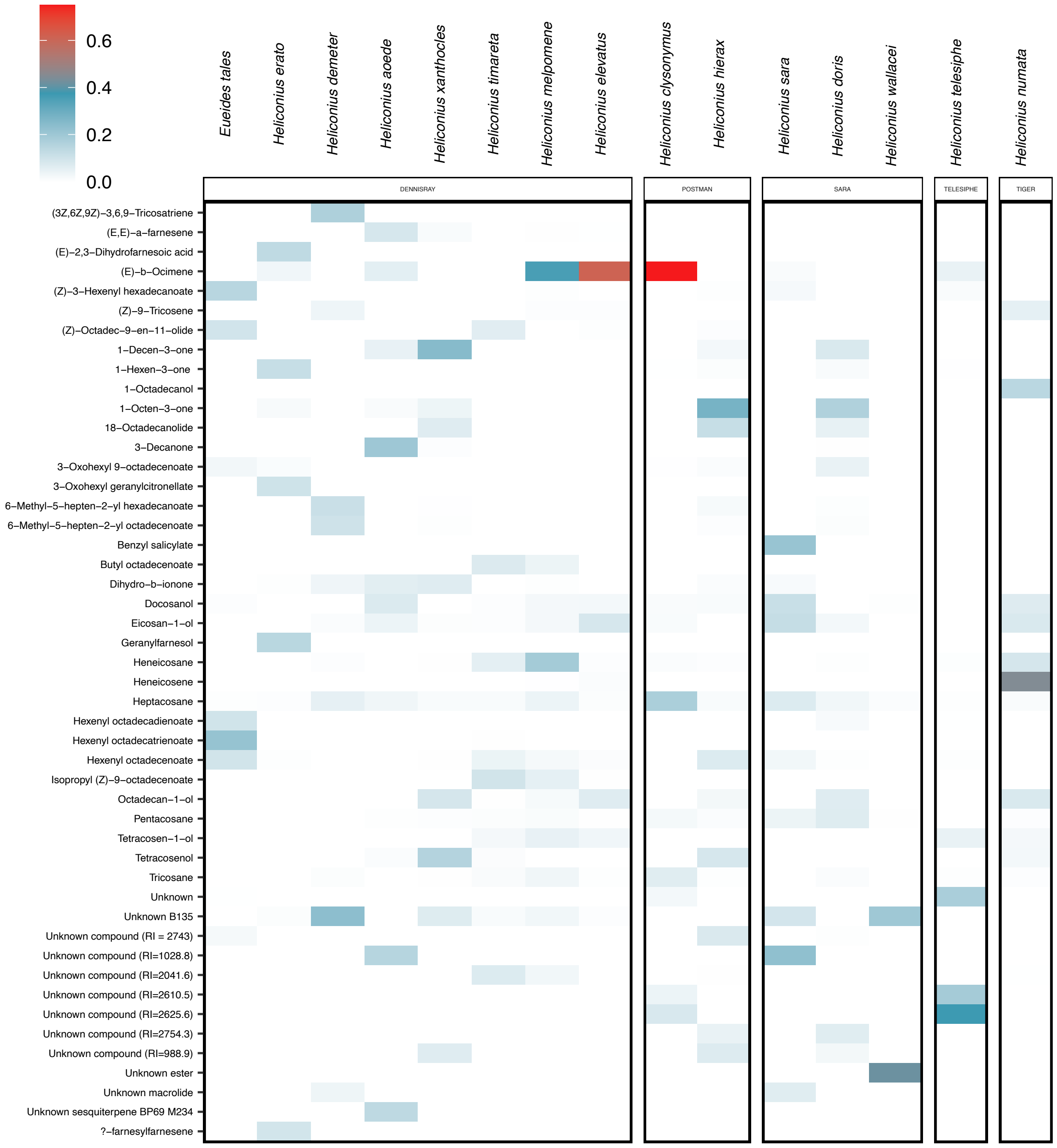


**Figure S4.** Variation in absolute compound concentration (nanograms) among the 50 most abundant Heliconiini clasper scent gland (CSG) compounds, with species grouped by mimicry.

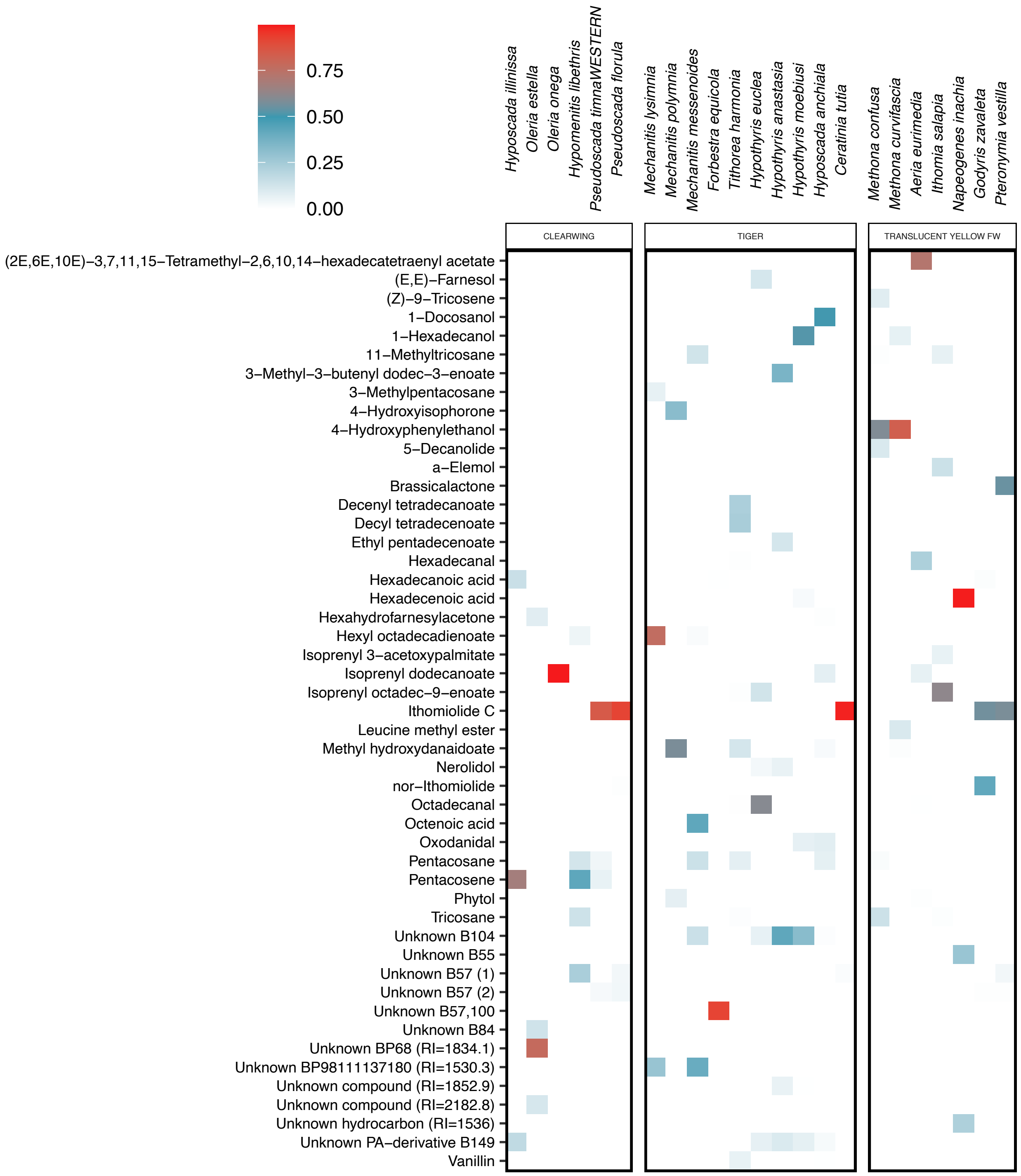


**Figure S5.** Variation in absolute compound concentration (nanograms) among the 50 most abundant Ithomiini androconial compounds, with species grouped by mimicry.

**Table S1.** Indicator androconial compounds significantly associated with Heliconiini species based on their specificity (occurrence in that species only) and fidelity (occurs in all members of that species). All compounds with a combined statistic score of 0.95 or over are included. Rows highlighted in grey represent species that do not have any compounds with a combined statistic score ≥ 0.95. For these species, the highest scoring indicator compound was selected instead.

| | | A | B | stat | *p* |
|---|---|---|---|---|---|
| ***Heliconius aoede*** | Hexadecanal | 1 | 1 | 1 | 0.001 |
| | Tetradecanoic acid | 1 | 1 | 1 | 0.001 |
| ***Heliconius clysonymus*** | Nonanoic acid | 1 | 1 | 1 | 0.001 |
| | Unknown compound (RI=1076.8) | 1 | 1 | 1 | 0.001 |
| | Unknown B148 | 0.98 | 1 | 0.99 | 0.001 |
| | (Z)-11-Eicosenol | 0.98 | 1 | 0.99 | 0.001 |
| ***Heliconius demeter*** | Tricosene | 1 | 1 | 1 | 0.001 |
| | Unknown branched alcohol | 1 | 1 | 1 | 0.001 |
| | (3Z,6Z,9Z)-3,6,9-Pentacosatriene | 1 | 1 | 1 | 0.001 |
| | Hexadecadienolide | 1 | 1 | 1 | 0.001 |
| | (3Z,6Z,9Z)-3,6,9-Tricosatriene | 1 | 1 | 1 | 0.001 |
| | (6Z,9Z)-6,9,Tricosadiene | 1 | 1 | 1 | 0.001 |
| | Phytol | 1 | 1 | 1 | 0.001 |
| | Unknown B43, 152 | 1 | 1 | 1 | 0.001 |
| | Docosadiene | 1 | 1 | 1 | 0.001 |
| | (3Z,6Z,9Z)-3,6,9-Docosatriene | 1 | 1 | 1 | 0.001 |
| | (Z)-9-Tricosene | 0.92 | 1 | 0.96 | 0.001 |
| ***Heliconius doris*** | Unknown B82,96 | 1 | 1 | 1 | 0.001 |
| ***Heliconius elevatus*** | Homovanillyl alcohol | 0.9 | 1 | 0.95 | 0.001 |
| ***Heliconius erato*** | T-Cadinol | 1 | 1 | 1 | 0.001 |
| ***Heliconius hierax*** | Unknown compound (RI=1493) | 1 | 1 | 1 | 0.002 |
| | Octadecanenitrile | 1 | 1 | 1 | 0.002 |
| | Eicosanenitrile | 1 | 1 | 1 | 0.002 |
| | Unknown B79 | 1 | 1 | 1 | 0.002 |
| | Icosylacetate | 0.96 | 1 | 0.98 | 0.002 |
| ***Heliconius melpomene*** | Octadecenolide | 1 | 1 | 1 | 0.001 |
| ***Heliconius numata*** | Heneicosane | 0.93 | 1 | 0.96 | 0.001 |
| ***Heliconius sara*** | Unknown BP99 | 0.93 | 1 | 0.97 | 0.001 |
| ***Eueides tales*** | 11-Methyltricosane | 0.78 | 1 | 0.88 | 0.01 |
| ***Heliconius telesiphe*** | Hexahydrofarnesyl acetone | 0.9 | 1 | 0.95 | 0.001 |
| ***Heliconius timareta*** | 5-Decanolide | 0.62 | 0.83 | 0.72 | 0.01 |
| ***Heliconius wallacei*** | Unknown compound (RI = 1514.6) | 1 | 1 | 1 | 0.001 |
| ***Heliconius xanthocles*** | Furanoid Linalooloxide | 1 | 1 | 1 | 0.001 |
| | Pyranoid linalooloxide | 1 | 1 | 1 | 0.001 |

**Table S2.** Indicator clasper scent gland compounds significantly associated with Heliconiini species based on their specificity (occurrence in that species only) and fidelity (occurs in all members of that species). All compounds with a combined statistic score of 0.95 or over are included. Rows highlighted in grey represent species that do not have any compounds with a combined statistic score ≥ 0.95. For these species, the highest scoring indicator compound was selected instead.

| | | **A** | **B** | **stat** | ***p*** |
|---|---|---|---|---|---|
| ***Heliconius aoede*** | Unknown Hydrocarbon BP97 | 1 | 1 | 1 | 0.001 |
| | RI.2830.6.49.0436min.HEA.001G | 1 | 1 | 1 | 0.001 |
| | Unknown Ester BP239.256 | 1 | 1 | 1 | 0.001 |
| | 3-Decanol | 1 | 1 | 1 | 0.001 |
| | 3-Dodecanone | 1 | 1 | 1 | 0.001 |
| | Sesquiterpene BP41,69 | 1 | 1 | 1 | 0.001 |
| | Unknown sesquiterpene BP69 M234 | 1 | 1 | 1 | 0.001 |
| ***Heliconius clysonymus*** | Unknown compound (RI=2887.8) | 1 | 1 | 1 | 0.001 |
| ***Heliconius demeter*** | Unknown compound (RI=1008.7) | 1 | 1 | 1 | 0.001 |
| | Unknown compound (RI=1094.1) | 1 | 1 | 1 | 0.001 |
| | Unknown compound (RI.2801.3) | 1 | 1 | 1 | 0.001 |
| | Unknown compound (RI=2869.2) | 1 | 1 | 1 | 0.001 |
| | Trans-epoxy-ocimene | 1 | 1 | 1 | 0.001 |
| | (3Z,6Z,9Z)-3,6,9-Tricosatriene | 1 | 1 | 1 | 0.001 |
| | (6Z,9Z)-6,9,Tricosadiene | 0.99 | 1 | 0.99 | 0.001 |
| ***Heliconius doris*** | RI.2563.5.45.1939min.HCL.11A | 1 | 1 | 1 | 0.001 |
| | RI.2682.5.46.9449min.HDE.003G | 1 | 1 | 1 | 0.001 |
| | Isoprenyloctadeca 2,11-dienoate | 1 | 1 | 1 | 0.001 |
| | Isoprenylpalmitate | 1 | 1 | 1 | 0.001 |
| | Unknown BP43 | 1 | 1 | 1 | 0.001 |
| | Unknown compound (RI=2330) | 1 | 1 | 1 | 0.001 |
| | Unknown compound (RI=2537.1) | 1 | 1 | 1 | 0.001 |
| | Unknown compound (RI=2769.6) | 1 | 1 | 1 | 0.001 |
| | Isoprenyloctadec-9-enoate | 0.91 | 1 | 0.96 | 0.001 |
| ***Heliconius elevatus*** | (*Z* )-beta-ocimene | 1 | 1 | 1 | 0.001 |
| | Sesquiterpene BP69 | 1 | 1 | 1 | 0.001 |
| ***Heliconius erato*** | 2-phenylethyloctadecanoate | 1 | 1 | 1 | 0.002 |
| | 2-nitroethyl-benzene | 1 | 1 | 1 | 0.002 |
| | (Z)-3-Hexenyl 3-methylbutyrate | 1 | 1 | 1 | 0.002 |
| | (Z)-3-Hexenyltiglate | 1 | 1 | 1 | 0.002 |
| | Hexyl-R(E)-2,3-dihydrofarnesenoate | 1 | 1 | 1 | 0.002 |
| | alpha-farnesylfarnesene | 1 | 1 | 1 | 0.002 |
| | alpha-geranylfarnesene | 1 | 1 | 1 | 0.002 |
| | beta-farnesylfarnesene | 1 | 1 | 1 | 0.002 |
| | beta-geranylfarnesene | 1 | 1 | 1 | 0.002 |
| | (2E,6E,10E)-3,7,11,15-Tetramethyl-2,6,10,14-hexadecatetraenylacetate | 1 | 1 | 1 | 0.002 |

| | | | | | |
|---|---|---|---|---|---|
| | (Z)-3,Hexenylgeranylcitronellate | 1 | 1 | 1 | 0.002 |
| | 2-Undecenylacetate | 1 | 1 | 1 | 0.002 |
| | 3-Nonanone | 1 | 1 | 1 | 0.002 |
| | 3-Oxoheptyl-(E)-2,3-dihydrofarnesenoate | 1 | 1 | 1 | 0.002 |
| | 3-Oxohexylgeranylcitronellate | 1 | 1 | 1 | 0.002 |
| | D-Limonene | 1 | 1 | 1 | 0.002 |
| | DHF ester | 1 | 1 | 1 | 0.002 |
| | Geranylcitronellicacid | 1 | 1 | 1 | 0.002 |
| | Geranylfarnesol | 1 | 1 | 1 | 0.002 |
| | Geranylgeranylacetone | 1 | 1 | 1 | 0.002 |
| | Hexyl3.methylbutyrate | 1 | 1 | 1 | 0.002 |
| | Hexylgeranylcitronellate | 1 | 1 | 1 | 0.002 |
| | Isoprenyl(E)-2.3.dihydrofarnesenoate | 1 | 1 | 1 | 0.002 |
| | Isoprenyl-Prenylgeranylcitronellate | 1 | 1 | 1 | 0.002 |
| | Mellein | 1 | 1 | 1 | 0.002 |
| | Unknown compound (RI=1770) | 1 | 1 | 1 | 0.002 |
| | Unknown compound (RI=2158.4) | 1 | 1 | 1 | 0.002 |
| | Sesterterpene (RI=2762) | 1 | 1 | 1 | 0.002 |
| | T-Cadinol | 1 | 1 | 1 | 0.002 |
| | Undecen-3-one | 1 | 1 | 1 | 0.002 |
| | Unknown BP71.117 | 1 | 1 | 1 | 0.002 |
| | 3-Oxooctyl S(E)-2,3,dihydrofarnesenoate | 0.92 | 1 | 0.96 | 0.002 |
| ***Heliconius hierax*** | Unknown BP110 (RI=2766.4) | 1 | 1 | 1 | 0.001 |
| | Unknown compound (RI=2554.5) | 1 | 1 | 1 | 0.001 |
| | Hexyloctadecanoate | 1 | 1 | 1 | 0.001 |
| ***Heliconius melpomene*** | E-E-E-alpha Springene | 1 | 0.75 | 0.87 | 0.001 |
| ***Heliconius numata*** | Eicosene | 1 | 1 | 1 | 0.001 |
| | TricoseneRI2286 | 0.97 | 1 | 0.99 | 0.001 |
| ***Heliconius sara*** | Benzoicacid-2-hydroxy phenylmethylester | 1 | 1 | 1 | 0.001 |
| ***Heliconius tales*** | (Z)-3-Hexenylheptadecanoate | 1 | 1 | 1 | 0.002 |
| | (Z)-3-Hexenylhexadecenoate | 1 | 1 | 1 | 0.002 |
| | 11-Methyltricosane | 1 | 1 | 1 | 0.002 |
| | 9-11-Hexadecadien-13-olide | 1 | 1 | 1 | 0.002 |
| | Hexadecatrienolide | 1 | 1 | 1 | 0.002 |
| | Hexenylester | 1 | 1 | 1 | 0.002 |
| | Hexyl 2-butenoate | 1 | 1 | 1 | 0.002 |
| | Hexyl hexadecadienoate | 1 | 1 | 1 | 0.002 |
| | Neroloxide related BP6885 | 1 | 1 | 1 | 0.002 |
| | Pentacosene | 1 | 1 | 1 | 0.002 |
| | Unknown B98 | 1 | 1 | 1 | 0.002 |
| | Octadecatrienoate | 0.99 | 1 | 1.00 | 0.001 |
| | Octadecadienoate ester | 0.89 | 1 | 0.95 | 0.004 |

| | | | | | |
|---|---|---|---|---|---|
| ***Heliconius telesiphe*** | Unknown compound (RI=1311.8) | 1 | 1 | 1 | 0.001 |
| | Unknown compound (RI=1615.7) | 1 | 1 | 1 | 0.001 |
| | Unknown compound (RI=2693.6) | 1 | 1 | 1 | 0.001 |
| | Unknown compound (RI=2722.9) | 1 | 1 | 1 | 0.001 |
| | 3-Oxohexyl (E)-2,3-dihydrofarneseoate | 0.99 | 1 | 1 | 0.001 |
| | 3,7-octadiene-2,6-diol-2,6-dimethyl | 0.93 | 1 | 0.97 | 0.001 |
| ***Heliconius timareta*** | Unknown compound (RI=2041.1) | 1 | 1 | 1 | 0.001 |
| | 2-sec-Butyl-3-methoxypyrazine | 1 | 1 | 1 | 0.001 |
| | 20-Eicosanolide | 1 | 1 | 1 | 0.001 |
| | Butyloctadecadienoate | 1 | 1 | 1 | 0.001 |
| | Icosenolide | 1 | 1 | 1 | 0.001 |
| | Isopropyloctadecadienoate | 1 | 1 | 1 | 0.001 |
| | Octadecadienolide | 1 | 1 | 1 | 0.001 |
| | Isopentyloctadecadecenoate | 1 | 1 | 1 | 0.001 |
| | Unknown compound (RI=2308) | 1 | 1 | 1 | 0.001 |
| | Octadecenolide | 0.98 | 1 | 0.99 | 0.001 |
| | Butyloctadecenoate | 0.93 | 1 | 0.97 | 0.001 |
| | (Z9,E11)-octadeca-9,11-dien-13-olide Isomer | 0.92 | 1 | 0.96 | 0.001 |
| | Unknown compound (RI=2366.1) | 0.91 | 1 | 0.96 | 0.001 |
| | Unknown compound (RI=2041.2) | 0.91 | 1 | 0.95 | 0.001 |
| | Ethyloleat | 0.89 | 1 | 0.95 | 0.001 |
| ***Heliconius wallacei*** | Unknown compound (RI=1959.7) | 1 | 1 | 1 | 0.001 |
| | UnknownBP43.138M.282.RI.1963.7.29.4100minS10523G | 1 | 1 | 1 | 0.001 |
| | Unknown BP57, 110 | 1 | 1 | 1 | 0.001 |
| | Unknown BP57, 96 | 1 | 1 | 1 | 0.001 |
| | Unknown BP57 | 1 | 1 | 1 | 0.001 |
| | Butyldecanoate | 1 | 1 | 1 | 0.001 |
| | Pyrimidin Derivative | 1 | 1 | 1 | 0.001 |
| | Unknown compound (RI=1841.8) | 1 | 1 | 1 | 0.001 |
| | Unknown compound (RI=1844.4) | 1 | 1 | 1 | 0.001 |
| ***Heliconius xanthocles*** | Unknown compound (RI=1383.2) | 1 | 1 | 1 | 0.001 |
| | Unknown compound (RI=2667.6) | 0.99 | 1 | 1 | 0.001 |

**Table S3.** Indicator androconial compounds significantly associated with Ithomiini species based on their specificity (occurrence in that species only) and fidelity (occurs in all members of that species). All compounds with a combined statistic score of 0.95 or over are included. Rows highlighted in grey represent species that do not have any compounds with a combined statistic score ≥ 0.95. For these species, the highest scoring indicator compound was selected instead.

| | | A | B | stat | *p* |
|---|---|---|---|---|---|
| ***Hypothyris anastasia*** | (E,E)-a-Farnesene | 1 | 1 | 1 | 0.0099 |
| | Unknown compound (RI=1882.9) | 1 | 1 | 1 | 0.0099 |
| ***Hyposcada anchiala*** | Pentacosene | 1 | 1 | 1 | 0.0099 |
| | 1-docosanol | 1 | 1 | 1 | 0.0099 |
| | 1-tricosanol | 1 | 1 | 1 | 0.0099 |
| ***Methona confusa*** | Heneicosane | 1 | 1 | 1 | 0.0099 |
| | (Z)-9-tricosene | 1 | 1 | 1 | 0.0099 |
| | 5-Decanolide | 1 | 1 | 1 | 0.0099 |
| | Unknown BP149,164 | 1 | 1 | 1 | 0.0099 |
| | Unknown compound (RI=2175.2) | 1 | 1 | 1 | 0.0099 |
| ***Methona curvifascia*** | Unknown compound (RI=1032) | 1 | 1 | 1 | 0.0099 |
| ***Forbestra equicola*** | Unknown B57,100 | 1 | 1 | 1 | 0.0198 |
| | Unknown B 132 | 1 | 1 | 1 | 0.0198 |
| | Edulan I | 1 | 1 | 1 | 0.0198 |
| | Unknown B100,118 | 1 | 1 | 1 | 0.0198 |
| | Hexadecanoic acid 2-hydroxyethyl amide | 1 | 1 | 1 | 0.0198 |
| | 2-Methylpentan-3-one | 1 | 1 | 1 | 0.0198 |
| | 3-Methylpent-3-en-2-ol | 1 | 1 | 1 | 0.0198 |
| ***Oleria estella*** | Unknown BP68 | 1 | 1 | 1 | 0.0099 |
| ***Hypothyris euclea*** | Unknown compound (RI=2189.3) | 1 | 1 | 1 | 0.0099 |
| | Octadecanal | 0.99 | 1 | 1 | 0.0099 |
| | Unknown compound (RI=2195.2) | 0.97 | 1 | 0.98 | 0.0099 |
| ***Aeria eurimedia*** | (2E,6E,10E)-3,7,11,15-Tetramethyl-2,6,10,14-hexadecatetraenylacetate | 1 | 1 | 1 | 0.0099 |
| | Hexadecanal | 0.97 | 1 | 0.99 | 0.0099 |
| ***Pseudoscada florula*** | Dihydroactinidiolide | 0.84 | 1 | 0.92 | 0.0198 |
| ***Tithorea harmonia*** | (Z)-3-Hexenylheptadecanoate | 1 | 1 | 1 | 0.0099 |
| | (Z)-3-Hexenyloctadecanoate | 1 | 1 | 1 | 0.0099 |
| | Decenyl Tetradecanoate | 1 | 1 | 1 | 0.0099 |
| | Decyl Tetradecenoate | 1 | 1 | 1 | 0.0099 |
| | Heptadecanal | 1 | 1 | 1 | 0.0099 |
| | Hexylhexadecenoate | 1 | 1 | 1 | 0.0099 |
| | Isopropyloctadecanoate | 1 | 1 | 1 | 0.0099 |
| | Mellein | 1 | 1 | 1 | 0.0099 |
| | Methyl farnesoate | 1 | 1 | 1 | 0.0099 |
| | Isopropyl octadecatrienoate | 1 | 1 | 1 | 0.0099 |
| | Isopropyl octadecenoate (2) | 1 | 1 | 1 | 0.0099 |
| | Hexyl hexadecenoate | 1 | 1 | 1 | 0.0099 |
| | Decenyl dodecanoate | 1 | 1 | 1 | 0.0099 |
| | Decyl dodecanoate + hexyl hexadecenoate (2) | 1 | 1 | 1 | 0.0099 |
| | Hexyl hexadecenoate | 1 | 1 | 1 | 0.0099 |
| | Hexyl octadecenoate | 1 | 1 | 1 | 0.0099 |
| | Octyl hexadecanoate | 1 | 1 | 1 | 0.0099 |
| | Unknown compound (RI=1398.3) | 0.98 | 1 | 0.99 | 0.0099 |

| | | | | | |
|---|---|---|---|---|---|
| ***Hyposcada illinissa*** | Hexadecanoic Acid | 0.92 | 1 | 0.96 | 0.0198 |
| ***Napeogenes inachia*** | Hydrocarbon (RI=1536.5) | 1 | 1 | 1 | 0.0099 |
| | Unknown B55 | 1 | 1 | 1 | 0.0099 |
| ***Hypomenitis libethris*** | 9-Tricosene | 1 | 1 | 1 | 0.0099 |
| | Tetracosane | 0.9 | 1 | 0.95 | 0.0099 |
| ***Mechanitis lysimnia*** | Unknown compound (RI=2533.1) | 0.93 | 1 | 0.96 | 0.0099 |
| ***Mechanitis messenoides*** | Ionone derivative BP125137167 | 1 | 1 | 1 | 0.0099 |
| ***Hypothyris moebuisi*** | 1-Hexadecanol | 0.92 | 1 | 0.96 | 0.0099 |
| ***Oleria onega*** | Unknown compound (RI=1854.5) | 0.92 | 1 | 0.96 | 0.0099 |
| ***Mechanitis polymnia*** | 4-hydroxyisophorone | 1 | 1 | 1 | 0.0198 |
| | Unknown branched alcohol (RI.1873.2) | 1 | 1 | 1 | 0.0198 |
| | Phytol | 0.94 | 1 | 0.97 | 0.0099 |
| ***Ithomia salapia*** | Isoprenyl (2E,11Z)-2,11-octadecadienoate | 1 | 1 | 1 | 0.0099 |
| | Isoprenyl (2E,13Z)-2,13-octadecadienoate | 1 | 1 | 1 | 0.0099 |
| | Isoprenyl 3-acetoxyhexadecanoate | 1 | 1 | 1 | 0.0099 |
| | Isoprenyl 3-acetoxyhexadecenoate | 1 | 1 | 1 | 0.0099 |
| | Isoprenyl octadecadienoate | 1 | 1 | 1 | 0.0099 |
| | Isoprenyl octadecatrienoate | 1 | 1 | 1 | 0.0099 |
| | RI,2523,4,44,9521minDSN436 | 1 | 1 | 1 | 0.0099 |
| | Unknown sesquiterpene | 1 | 1 | 1 | 0.0099 |
| | a-Elemol | 1 | 1 | 1 | 0.0099 |
| ***Pseudoscada timnaWESTERN*** | 11-Methylpentacosane | 1 | 1 | 1 | 0.0099 |
| ***Ceratinia tutia*** | Hydrocarbon (RI=1758.8) | 1 | 1 | 1 | 0.0297 |
| ***Pteronymia vestilla*** | Brassicalactone | 1 | 1 | 1 | 0.0099 |
| | Unknown compound (RI=1042.2) | 1 | 1 | 1 | 0.0099 |
| ***Godyris zavaleta*** | Unknown compound (RI=1194.1) | 0.99 | 1 | 1 | 0.0099 |

**Table S4.** Primers used for targeting and amplifying COI region of Heliconiini and Ithomiini butterflies.

| Sequence Name | Primer Sequence | Adapter |
|---|---|---|
| IAW_F1_Adapt | TGG AAT TTG AGC AGG AAT AGT AGG | TCG TCG GCA GCG TCA GAT GTG TAT AAG AGA CAG |
| IAW_R1_Adapt | AA TAG CTA AAT CAA CAG AAG AAC CT | GTC TCG TGG GCT CGG AGA TGT GTA TAA GAG ACA G |
